# Kinetochore-independent mechanisms of sister chromosome separation

**DOI:** 10.1101/2020.08.04.236943

**Authors:** Hannah Vicars, Travis Karg, Brandt Warecki, Ian Bast, William Sullivan

## Abstract

Although kinetochores normally play a key role in sister chromatid separation and segregation, chromosome fragments lacking kinetochores (acentrics) can in some cases separate and segregate successfully. In *Drosophila* neuroblasts, acentric chromosomes undergo delayed, but otherwise normal sister separation, revealing the existence of kinetochore-independent mechanisms driving sister chromosome separation. Bulk cohesin removal from the acentric is not delayed, suggesting factors other than cohesin are responsible for the delay in acentric sister separation. In contrast to intact kinetochore-bearing chromosomes, we discovered that acentrics align parallel as well as perpendicular to the mitotic spindle. In addition, sister acentrics undergo unconventional patterns of separation. For example, rather than the simultaneous separation of sisters, acentrics oriented parallel to the spindle often slide past one another toward opposing poles. To identify the mechanisms driving acentric separation, we screened 117 RNAi gene knockdowns for synthetic lethality with acentric chromosome fragments. In addition to well-established DNA repair and checkpoint mutants, this candidate screen identified synthetic lethality with X-chromosome-derived acentric fragments in knockdowns of Greatwall (cell cycle kinase), EB1 (microtubule plus-end tracking protein), and Map205 (microtubule-stabilizing protein). Additional image-based screening revealed that reductions in Topoisomerase II levels disrupted sister acentric separation. Intriguingly, live imaging revealed that knockdowns of EB1, Map205, and Greatwall preferentially disrupted the sliding mode of sister acentric separation. Based on our analysis of EB1 localization and knockdown phenotypes, we propose that in the absence of a kinetochore, microtubule plus-end dynamics provide the force to resolve DNA catenations required for sister separation.

**AUTHOR SUMMARY:** Kinetochores, the site on the chromosomes to which microtubules attach driving the separation and segregation of replicated sister chromosomes, have been viewed as essential for proper cell division and accurate transmission of chromosomes into daughter cells. However previous studies demonstrated that sister chromosomes lacking kinetochores (acentrics) often undergo separation, segregation and transmission. Here we demonstrate that sister acentrics are held together through DNA intertwining. We show that during anaphase, acentric sister separation is achieved through Topoisomerase activity, an enzyme that resolves these DNA linkages, as well as forces generated on the acentrics by the growing ends of highly dynamic microtubule polymers. We found that acentric sister chromatids display unique patterns of separation using mechanisms independent of the kinetochore. Additionally, we identified the specific microtubule-associated proteins required for the successful mitotic transmission of acentric chromosomes to daughter cells. These studies reveal unsuspected, distinct forces that likely act on all chromosomes during mitosis independent of kinetochore-microtubule attachments.

## INTRODUCTION

Eukaryotic cells have evolved mechanisms to detect and protect against genomic insults. These mechanisms include checkpoint pathways that delay cell cycle progression allowing time for repair as well as apoptotic pathways that eliminate the damaged cells from the dividing population [1]. Although a great deal is known regarding the function of these corrective pathways during interphase, much less is known about the mechanisms that protect against genomic instability after exit from metaphase. Studies demonstrate that DNA damage persisting through metaphase delays anaphase onset. This delay is mediated both by the DNA damage and spindle assembly checkpoint pathways [2, 3].

Despite these mechanisms, if the DNA damage remains, the checkpoints are overridden, and the cell exits metaphase [4]. The persistence of unrepaired double-strand breaks (DSBs) at metaphase is particularly problematic due to the formation of chromosome fragments, one of which lacks a telomere and the other lacking a kinetochore and a telomere. The latter type are known as acentrics and are incapable of forming canonical microtubule-kinetochore attachments that drive sister chromosome separation and segregation. Consequently, acentrics would be expected to lag on the metaphase plate and exhibit severe segregation defects. In accord with this expectation, acentrics often fail to segregate, are excluded from daughter nuclei, and subsequently form cytoplasmic micronuclei [5–7]. However, a growing number of reports demonstrate poleward migration of acentric chromosome fragments [5, 8–13]. Proposed mechanisms of acentric segregation include neo-centromere formation [14] and direct association of the acentric chromosome with microtubules [15] or a kinetochore-bearing chromosome [16–18].

Acentrics are efficiently induced in *Drosophila* bearing an I-CreI endonuclease transgene, which fortuitously recognizes a repetitive sequence within the pericentric rDNA repeats of the *Drosophila* X chromosome [19–22]. Induction of I-CreI expression results in the formation of acentrics in over 80% of third instar larval neuroblast cells [12]. Although acentrics lag behind on the metaphase plate well after the intact chromosomes migrate toward opposite poles, the acentrics have a remarkable ability to accurately separate, segregate and incorporate into daughter telophase nuclei [23]. A previous study done in the lab found that acentric segregation relies on the chromokinesin Klp3A and interpolar microtubules [15]. However, it remains unclear how sister acentrics are held together on the metaphase plate well after the main chromosome mass has separated. Additionally, it is unknown how acentric sisters are able to initially separate from one another instead of segregating together poleward. These behaviors reveal the existence of kinetochore-independent mechanisms maintaining sister chromosome association on the metaphase plate and driving their separation during anaphase. Possible explanations include delayed acentric cohesin removal, delayed resolution of sister DNA catenations, or opposing plus-end directed microtubule forces acting on each sister. Here we employ a combination of synthetic lethal screens and live imaging to identify factors required for proper separation of sister acentrics. Live analysis reveals three distinct modes of acentric separation: unzipping, sliding, and simultaneous dissociation. This candidate screen revealed that Topoisomerase II, the cell-cycle regulator Greatwall kinase, the microtubule (MT) plus-end tracking protein EB1, and the MT-associated protein Map205 provide key roles in sister separation of acentrics. In addition, gene knockdowns of EB1, Map205, and Greatwall preferentially disrupt the sliding mode of sister separation. As will be discussed, this analysis demonstrates the existence of kinetochore-independent mechanisms facilitating sister chromosome separation.

## RESULTS

### Acentric sister separation, but not cohesin removal, is delayed during the metaphase-to-anaphase transition

As described previously, acentric chromosome fragments are efficiently generated through heat-shock induction of an I-CreI transgene that specifically targets and creates DSBs in the rDNA repeats at the base of the *Drosophila* X chromosome [19]. In accord with previous studies [12], live analysis of the resulting X chromosome acentrics in the larval neuroblasts reveals that sister separation of the acentrics occurs on average 148 seconds (± 44, N=19) after separation of the intact chromosomes (**Figure 1**). We define acentric sister separation as the point in which the sister acentrics can be clearly distinguished. Acentric sister segregation is defined as the interval between separation of sisters and their migration to the spindle pole. The delay in timing of acentric separation is defined as the time elapsed between the initiation of intact sister chromosome separation and that of acentric sister separation. Sister chromatids are held together at metaphase by cohesin, a tripartite ring-like protein complex comprised of two structural maintenance of chromosome proteins (SMC1, SMC3) and a kleisin subunit (Rad21/Scc1) [24, 25]. Initially, cohesin removal occurs only along the chromosome arms through proteolytic cleavage of the kleisin subunit by separase just prior to anaphase onset [26–28]. Due to the Sgo/PP2A-dependent protection mechanism, cohesin remains at the centromeric regions [29]. Once cohesin is removed, microtubules drive sister separation [30, 31].

**Fig 1:**
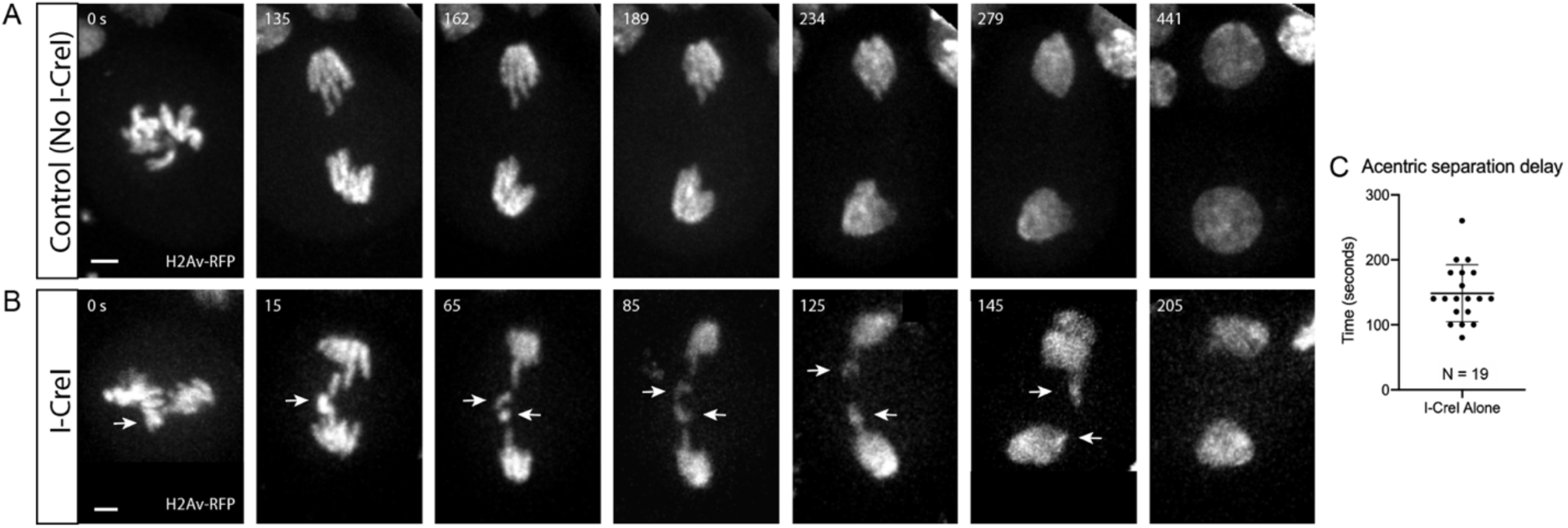
Acentric sister separation is delayed relative to kinetochore-bearing chromosomes during anaphase. (A) Still frames of a time-lapse movie of a mitotic neuroblast labeled with H2Av-RFP and not expressing I-CreI. (B) Still frames of a time-lapse movie of a mitotic neuroblast with I-CreI induced acentrics. Separation of sister acentrics (arrows) is delayed. Consequently, they lag on the spindle equator but eventually separate, segregate, and are reincorporated into daughter nuclei. Bars, 2 μm. Time in seconds. (C) Scatterplot showing the delay in acentric sister separation after anaphase onset. Delay (seconds) was measured from when kinetochore-bearing chromosomes initiated separation to when acentric sister chromosomes initiated separation. Bars represent mean and standard deviations.

To investigate if delays in cohesin release are responsible for the delayed separation of acentric chromosomes, female neuroblast divisions were live imaged with a cohesin component, Rad21, tagged with EGFP [32]. Time-lapse images of a control neuroblast expressing Rad21-EGFP are shown in **Figure 2A** (chromosomes in magenta, cohesin in green). No lagging chromosomes are observed and Rad21 is cleared off of all chromosomes just prior to anaphase onset and separation of sister chromatids. **Figure 2B** and **Movie S1** show time-lapse images of an I-CreI-expressing neuroblast division. Separation and segregation of sister acentrics is delayed relative to the intact chromosomes. Interestingly, cohesin removal just prior to anaphase onset from the acentrics and the intact kinetochore-bearing chromosomes occurs simultaneously. This is more clearly seen in the single channel black and white cohesin images of **Figure 2B** (depicting yellow outlined regions of **Figure 2B**). Quantification of the Rad21 fluorescent signal supports the conclusion of a relatively synchronous removal of cohesin on the acentric and intact chromosomes (**Figure 2C**). The finding that sister acentrics remain paired despite the absence of cohesin and well after the intact chromosomes have separated, indicate additional forces must hold sister acentrics together.

**Fig 2:**
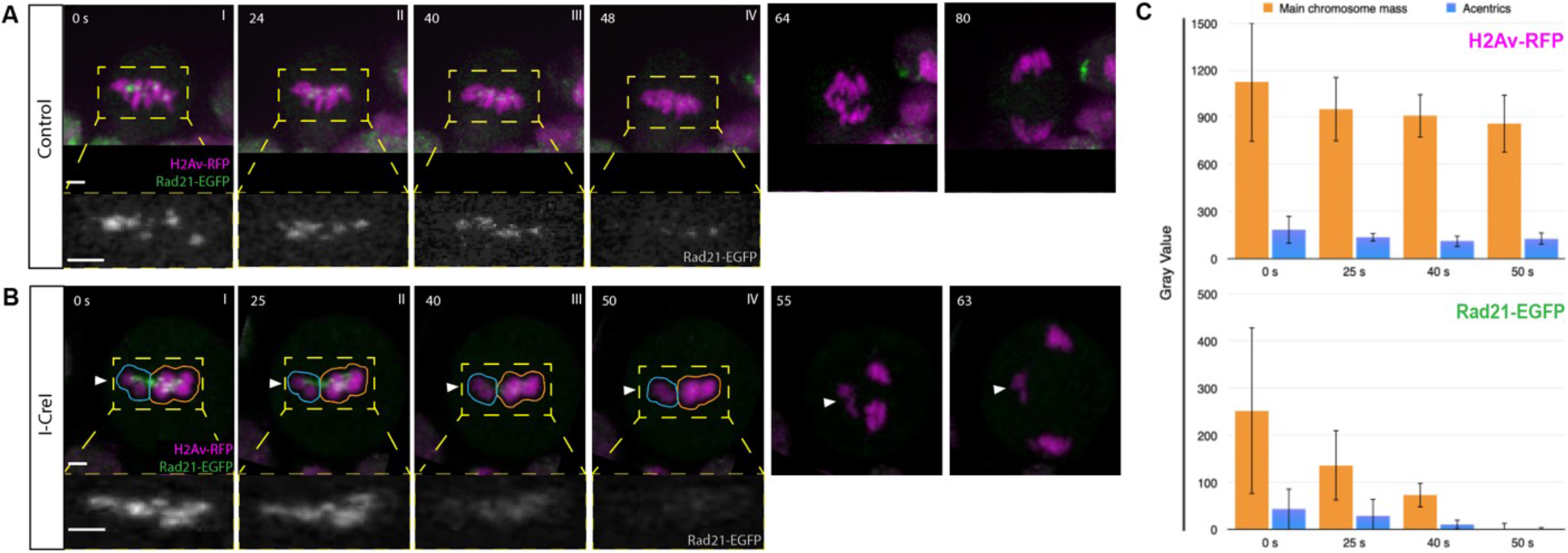
Cohesin complexes are cleared off of acentric sisters upon anaphase onset. Chromosomes labeled with H2Av-RFP (magenta) and cohesin labeled with Rad21-EGFP (green). (A) Stills from a time-lapse movie of a control mitotic neuroblast. (B) Still images from a time-lapse movie of a mitotic neuroblast with I-CreI-induced acentrics (Movie S1). Acentrics (arrowhead) lag on the spindle equator. (C) Bar graphs of a compilation of five videos of I-CreI-expressing neuroblasts showing the relative fluorescence intensities in arbitrary units (AU) of chromosomes (H2Av-RFP, top) cohesin (Rad21-EGFP, bottom) around acentrics (cyan outlined region) and the main mass of chromosomes (yellow outlined region) at time points 50, 40, 25, 0 s, prior to anaphase onset, respectively. Bars, 2 μm. Time in seconds. Error bars represent standard deviations of fluorescence intensities at all points tested.

### Acentric sister separation occurs via three distinct patterns

To examine dynamics of acentric separation, we imaged live neuroblasts expressing CreI, the histone marker H2Av-RFP, and the telomere marker HOAP-GFP [33]. Marked telomeres enabled us to determine the orientation of acentrics with respect to one another as they aligned on the metaphase plate, separated, and segregated poleward [15]. We observed three distinct patterns of sister acentric separation (**Figure 3**). In the most frequent pattern (49%, N=45), acentric pairs separate by sliding past one another (**Figure 3A**, Top row: histone-labeled chromosomes in magenta, HOAP-labeled telomeres in green). This is more clearly observed in the single histone channel movie (**Figure 3A** Bottom row and **Movie S2**). Also, in contrast to intact kinetochore-bearing chromosomes which always align perpendicular to the spindle, the paired sister acentrics align either parallel or perpendicular to the spindle and division axis.

**Fig 3:**
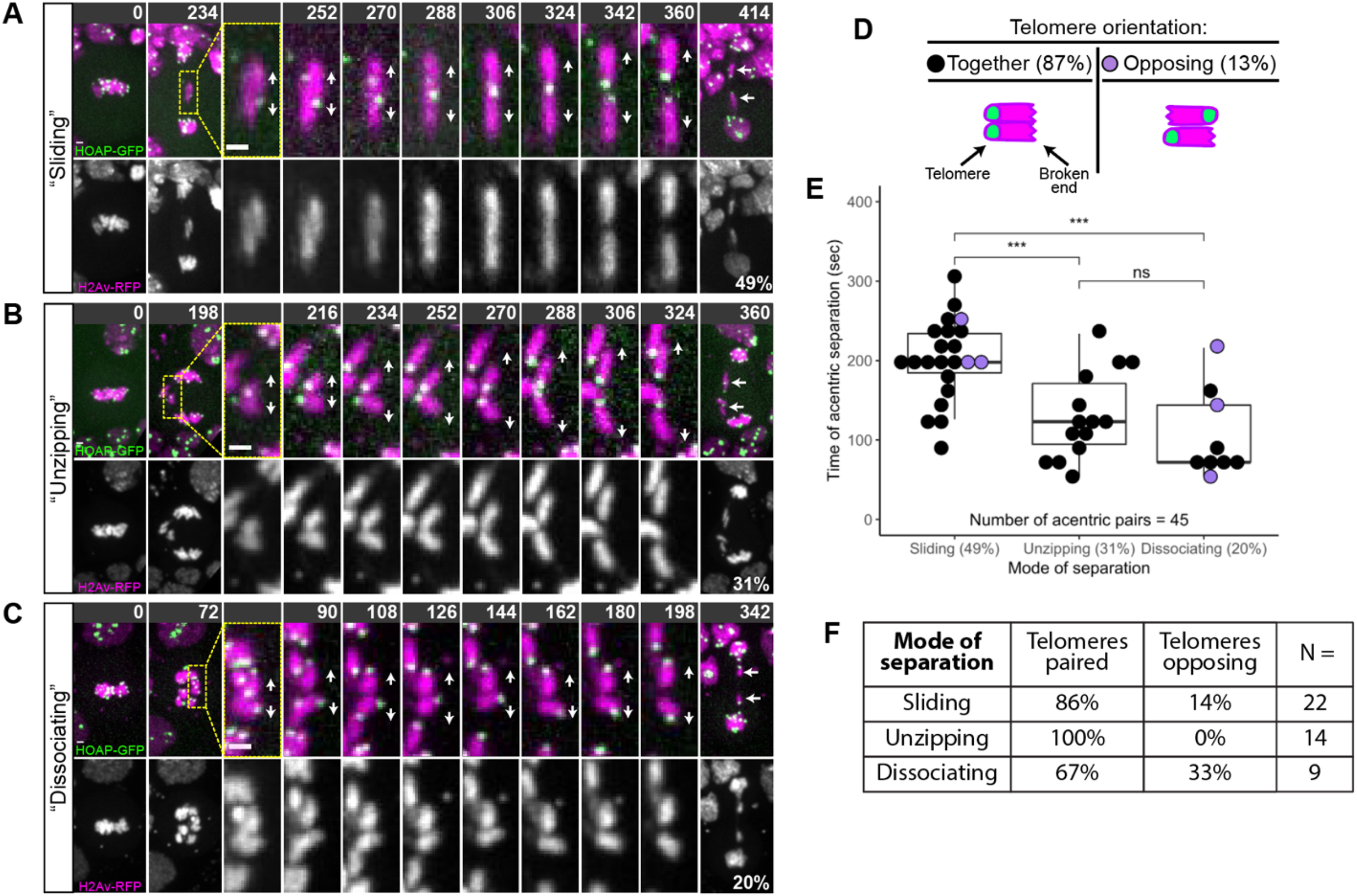
Sister acentrics separate via three distinct patterns. Chromosomes labeled with H2Av-RFP (magenta) and telomeres labeled with HOAP-GFP (green). Telomeres indicated by yellow arrowheads. Direction of separation indicated by white arrows. (A) Top row: Still images of a time-lapse movie of a mitotic neuroblast with I-CreI induced acentrics showing sister acentrics (cyan arrows) lagging behind during anaphase, paired with telomeres opposing, sliding past one another, and ultimately separating. Bottom row: Black and white images of sister acentrics sliding past one another (see arrows) (Movie S2). (B) Top row: Still images of a time-lapse movie of a mitotic neuroblast with I-CreI induced acentrics showing acentrics (cyan arrows) lagging on the metaphase plate. In a process we term “unzipping”, sister acentrics are aligned with telomeres paired, initiate separation at their broken ends followed by separation of telomeres. Bottom row: Black and white images of sister acentrics unzipping (see arrows). (C) Top row: Still images of a time-lapse movie of a mitotic neuroblast with I-CreI induced acentrics showing sister acentrics (cyan arrows) lagging behind during anaphase. In a process we term “simultaneous dissociation”, sister acentrics simultaneously separate along their entire length. Bottom row: Black and white images of sister acentrics simultaneously separating along their entire length (see arrows). (D) Frequency of acentric sisters paired with their telomeres aligned or in opposite orientations. (E) Measurements of the frequency of each mode of acentric sister separation (x-axis) and the timing of acentric sister separation after intact chromosomes separate (y-axis). Each dot represents on acentric pair. Black dots represent acentric sisters with telomeres aligned and purple dots represent acentric sisters with telomeres oriented in opposite directions. Boxes show interquartile ranges and lines show medians of the measured data. Asterisks represent statistical significance (***P=0.0008) determined by Mann-Whitney tests. (F) Chart showing the distribution of telomere orientations within each mode of acentric sister separation. Acentrics that separate by unzipping are always oriented with their telomeres paired.

The second pattern of acentric separation occurs via an “unzipping” mechanism (**Figure 3B**). This pattern occurs at a frequency of 31% (N=45). During separation, sister acentrics often first separate at the broken end followed by separation at the telomere. This is illustrated in **Figure 3B**: the telomeres remain associated while the broken ends are well separated. Rarely, the paired acentrics unzip from the telomere end first. In contrast to the acentrics that separate by sliding, these paired sister acentrics are frequently aligned perpendicular to the spindle and division axis.

In the third pattern of separation, the remaining 20% (N=45), acentric sisters cleanly separate from one another along their entire length similar to that observed for centric chromosomes (**Figure 3C**). We termed this centric-like pattern: simultaneous dissociation. Acentrics that separate via simultaneous dissociation are aligned in multiple orientations on the metaphase plate (from parallel to perpendicular) with respect to the spindle and division axis. At separation, dissociating sister acentrics simultaneously separate along their entire lengths.

We next analyzed the orientation of sister acentrics with respect to one another at the time of separation through labeling telomeres with HOAP-GFP. As expected, the vast majority (87%, 39/45) of sister acentrics were oriented with their telomeres paired and aligned (**Figure 3D**). However, a small but notable fraction (13%, 6/45), aligned with their telomeres opposed (**Figure 3D**). Acentric pairs that align with telomeres opposed presumably have already lost catenation and/or cohesin in order to adopt this geometry. 14% and 33% of acentric sisters that separated by sliding and dissociating, respectively, aligned with their telomeres opposed. Interestingly, sister acentrics that separated by unzipping were never observed aligned with their telomeres opposed. This suggests that there is an absolute requirement for sister pairing in acentrics that separate by unzipping.

To further characterize these three modes of acentric separation, we measured the time from anaphase onset (as determined by separation of the intact chromosomes) to sister acentric separation (**Figure 3E**). Sliding acentrics separated much later than acentrics (202 ± 51 seconds) that separated either by unzipping (131 ± 54 seconds) or dissociating (106 ± 55 seconds) (N=45, **Figure 3E**). These differences were statistically significant as determined by two-sided Mann-Whitney tests (P=0.0008 and P=0.0008, respectively). We did not detect any difference in the timing of acentric sister separation by unzipping or dissociating (P=0.25 as determined by a two-sided Mann-Whitney test).

Taken together, these results demonstrate three distinct patterns for acentric separation. We note that based on the movements and orientation of both sliding and dissociating acentrics, it is possible that catenations are lost along the entire length of the acentric pair simultaneously at the moment of separation. In contrast, acentrics that separate by unzipping and appear to have an absolute requirement for sister pairing may be more tightly associated with one another, and this tight association may be due to lingering catenations.

### Synthetic lethal screen identifies genes required for separation of sister acentrics

To identify the mechanisms required for transmission of acentric sister chromosomes, we screened candidate RNAi gene knockdowns that resulted in synthetic lethality in the presence of acentrics. The rationale for this screen is based on previous studies demonstrating that I-CreI induction of acentric chromosomes during third instar larval stage resulted in only slight reductions in adult survival because the acentrics are efficiently transmitted to daughter nuclei [12]. However, reducing or partially disrupting the function of genes required for the normal acentric transmission results in a dramatic reduction in adult survival upon acentric induction [12, 15]. Thus, we expected that a subset of the gene knockdowns that resulted in synthetic lethality upon I-CreI induction would be required for proper pairing and segregation of sister acentrics.

To perform the screen, adult flies bearing a heat-shock inducible I-CreI endonuclease and Gal4 under the control of a ubiquitously expressed Actin enhancer element (Act5) were crossed to adults bearing UAS-gene specific RNAi constructs. Lethality of heat-shocked (I-CreI induced) and non-heat-shocked (I-CreI not induced) F1 progeny bearing both constructs were assayed. RNAi constructs that significantly increased lethality upon I-CreI induction were of particular interest. We screened 117 candidate genes for synthetic lethality upon acentric induction (**Table S1**). These included genes encoding proteins spanning a diversity of mitotic functions, including microtubule-associated proteins, chromatin remodelers, DNA repair genes, cell cycle kinases, and cell cycle checkpoints. For each RNAi line, we determined the survival ratio of the RNAi knockdown with I-CreI induced to the RNAi knockdown alone (RNAi knockdown + I-CreI/RNAi knockdown) (**Table 1**). Because these RNAi knockdowns do not cause complete lethality, we classified hits as having a survival ratio of 65% or less (**Table 1**).

**Table 1:**
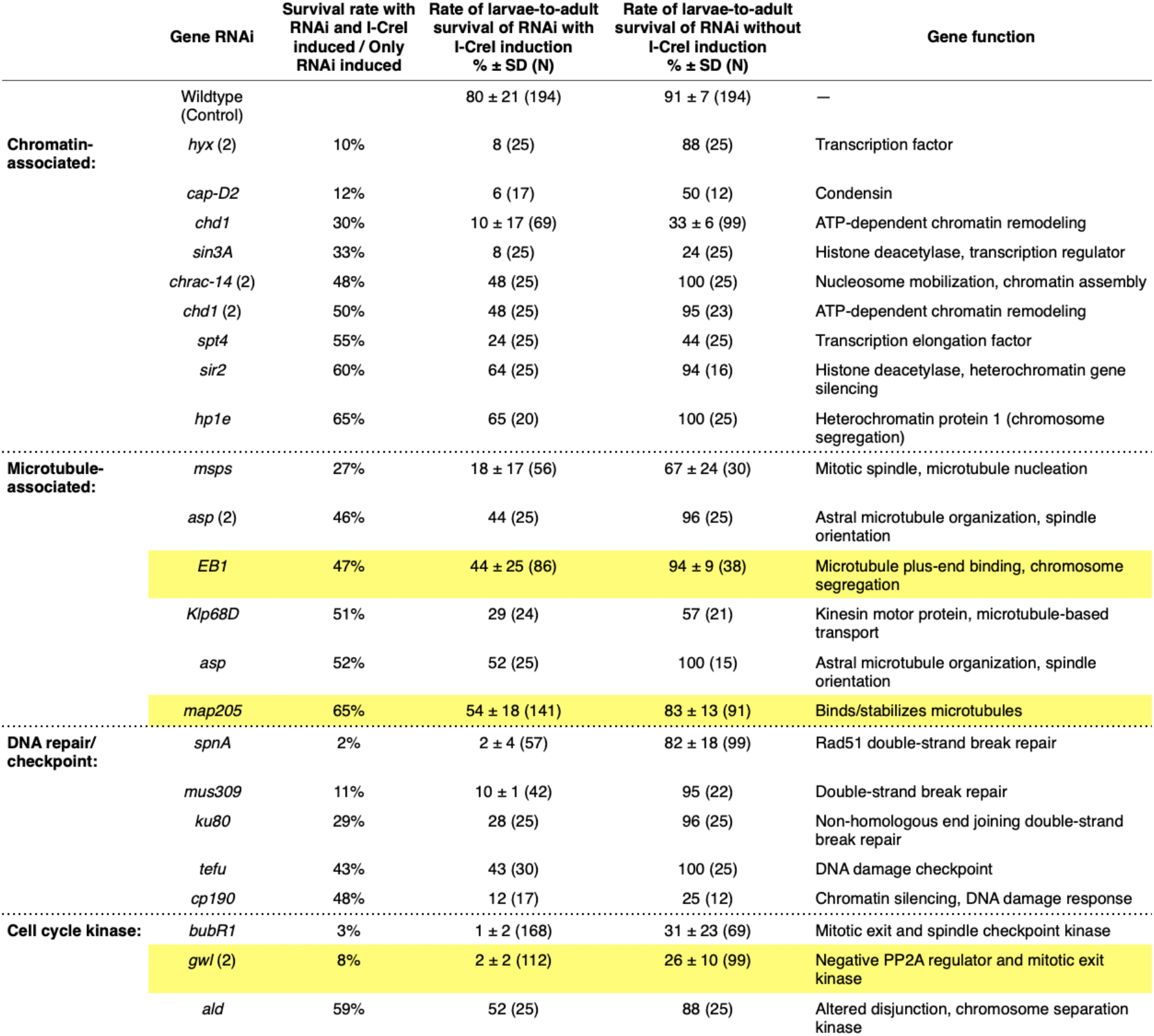
Top hits from synthetic lethality screen. Top hits from the synthetic lethality screen were grouped by gene function. Overall survival rate ratio was determined by the following: (percentage of surviving larvae after RNAi and I-CreI induction) / (percentage of surviving larvae after only RNAi induction). Those RNAi transgenes that resulted significant reduction in survival (a ratio of less than 0.65) were considered for follow-up live analysis.

The acentrics are generated through I-CreI induced double-strand breaks. As expected, RNAi knockdowns of DNA repair and cell cycle checkpoints exhibit synthetic lethality upon I-CreI induction. Genes involved in DNA repair all exhibit strong synthetic lethality upon I-CreI induction. DSB repair genes *spnA/Rad51* [34], *mus309* [35] and NHEJ gene *Ku80* [36] exhibited survival ratios of 2%, 11%, and 29%, respectively (**Table 1**). The DNA damage checkpoint gene *tefu* (ATM [37]) also results in a pronounced synthetic lethality (survival ratio of 43%). The screen also yielded a number of microtubule-associated proteins. These included *msps* (microtubule nucleation [38]), *asp* (astral microtubule organization [39]), *eb1* (plus-end microtubule binding [40]), *klp68D* (kinesin motor protein [41]), and *map205* (microtubule stabilizer [42]). Given that microtubule-based transport plays a key role in the poleward transport of the acentric chromosome fragments [15], synthetic lethal interactions with microtubule-associated proteins were expected. However, it was unclear whether microtubules and their associated proteins also play a role in the initial separation of sister acentrics during the metaphase-to-anaphase transition. The screen also yielded a large class of genes involved in chromatin organization. Synthetic lethal interactions with chromatin organizing proteins were expected due to the presence of I-CreI induced DSBs. This included *cap-D2* (condensin [43]), *chd1* (ATP-dependent chromatin remodeler [44]), *sin3A* and *sir2* (histone deacetylases [45]), *chrac* (nucleosome mobilization [46]), and *hp1* (heterochromatin protein [47]). Whether these are directly required for double-strand break repair, separation and/or segregation of sister acentrics remains to be determined. In accord with previous work, reduced levels of BubR1 kinase (spindle-assembly checkpoint protein) also exhibited a pronounced synthetic lethality upon I-CreI induction [12]. The screen also yielded two additional cell cycle kinases: *gwl* (greatwall kinase, an inhibitor of the cell cycle phosphatase PP2A [48]) and *ald* (altered disjunction, chromosome segregation [49]). We chose to focus on EB1 and Map205 due to their well-documented association with microtubules [40, 42]. Greatwall was chosen for follow-up because previous studies demonstrated that this kinase is required for sister separation of intact chromosomes [50].

### Live imaging analysis reveals the microtubule-associated proteins Map205 and EB1 and Topo II are required for separation, but not segregation of sister acentrics

We conducted live imaging experiments on neuroblasts to investigate the effect of specific RNAi-mediated gene knockdowns on acentric mitotic transmission. Each line contains an RFP-tagged histone transgene facilitating live confocal analysis [51]. Based on previous studies revealing the role of microtubules in acentric transmission [15], we initially focused on the microtubule-stabilizing protein Map205 and the microtubule plus-end associated protein EB1.

In a wild-type background, the majority of acentrics line up at the outer edge of the metaphase plate, separated from the main mass of intact chromosomes [12, 15]. As described above and in previous publications, during anaphase sister acentrics remain paired on the metaphase plate well after the separation of intact chromosomes (**Figure 4A**) [12, 15]. On average, separation of sister acentrics occurs 148 seconds ± 44 (N=19) after separation of the kinetochore-bearing chromosomes (**Table 2**). 83% (19/23) of acentric sister chromatid pairs separate normally with sister acentrics going to opposite cell poles (**Figure 4C**). In the remaining 17% (4/23), acentric sister chromatids line up in metaphase, lag behind at the metaphase plate, and segregate together to one pole of the cell (**Table 3**). Micronuclei form in telophase in 24% (4/17) of neuroblasts expressing I-CreI micronuclei form in telophase (**Figure 4D**).

**Fig 4:**
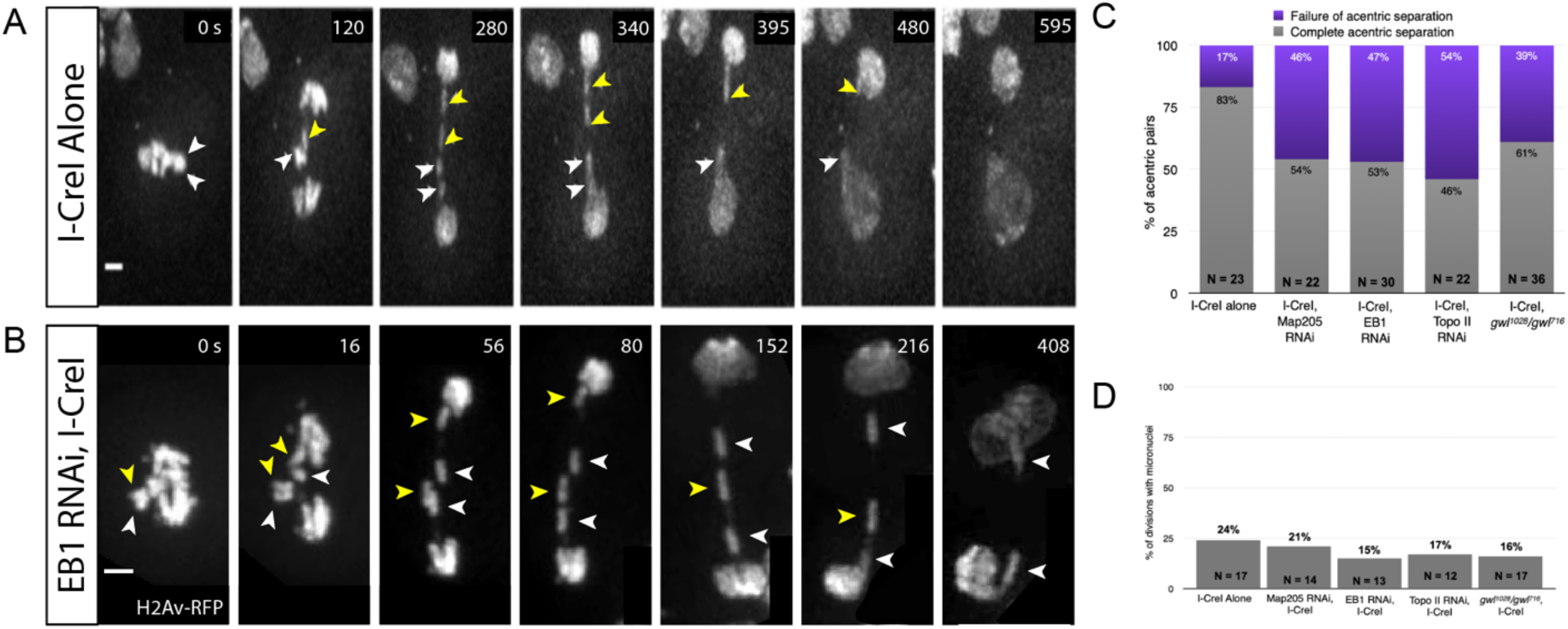
EB1 RNAi knockdowns specifically disrupt acentric sister separation. (A) Still frames of a time-lapse movie of a mitotic neuroblast with I-CreI induced acentrics. Paired sister acentrics (white and yellow arrowheads) lag behind at the spindle equator but eventually separate, segregate, and are reincorporated into daughter nuclei. (B) Still frames of a time-lapse movie of a mitotic neuroblast with I-CreI induced acentrics and expressing EB1 RNAi. Separation of paired sister acentrics (white and yellow arrowheads) fails, resulting in sisters segregating to and incorporating into the same daughter nucleus. Bars, 2 μm. Time in seconds. (Video 3) (C) Percentages of neuroblast divisions in which acentric sisters failed to completely separate from one another. (D) Percentages of neuroblast divisions in which acentrics failed to incorporate into daughter nuclei and formed one or more micronuclei.

**Table 2:** Sister separation of acentrics is delayed relative to sister separation of intact chromosomes. Average time (in seconds) for sister acentrics to initiate separation after sister kinetochore-bearing chromosomes separated. Time (average ± SD) measured from initiation of intact chromosome separation to initiation of acentric sister chromosome separation. Asterisks indicate statistical significance (**P=0.001, *P=0.003) as determined by two-sided Mann-Whitney tests.

| | Avg ± SD<br>(seconds) | N<br>(acentric pairs) |
| --- | --- | --- |
| **Control** | 148 ± 44 | 19 |
| **MAP205 RNAi** | 97 ± 39 ** | 12 |
| **EB1 RNAi** | 142 ± 57 | 16 |
| **Topo II RNAi** | 226 ± 77 * | 10 |
| ***gwl<sup>1028</sup>/gwl<sup>716</sup>*** | 138 ± 49 | 22 |

**Table 3:** Acentric separation occurs through three distinct modes. Acentric sister chromatids either fail to separate or separate from one another by three different modes: laterally sliding past one another, unzipping from one another, or simultaneously and evenly dissociating along their lengths.

|  | Fails to separate | Dissociate | Slides Apart | Unzip | N<br>(acentric pairs) |
| --- | --- | --- | --- | --- | --- |
| **Control** | 17% | 13% | 48% | 22% | 23 |
| **MAP205 RNAi** | 46% | 9% | 9% | 36% | 22 |
| **EB1 RNAi** | 47% | 20% | 13% | 20% | 30 |
| **Topo II RNAi** | 54% | 14% | 27% | 5% | 22 |
| ***gwl<sup>1028</sup>/gwl<sup>716</sup>*** | 39% | 11% | 28% | 22% | 36 |

Live imaging of acentric behavior in neuroblasts expressing Map205 RNAi revealed that 46% (10/22) of acentric sister chromatids do not separate from one another and segregate to one cell pole together, in comparison to 17% (4/23) in a wildtype background (**Figure 4C, Figure 5B, Table 3**). Additionally, after partial knockdown of Map205, sister acentrics separated from one another significantly earlier than acentrics in a wildtype background (P=0.001, Mann-Whitney test, **Table 2**). In spite of defects in acentric sister chromatid separation, their average poleward segregation velocity was normal (**Table 4**). Despite the increase in failed acentric separation, there was not an increase in micronuclei formation (21% compared to 24% in controls) (**Figure 4D**).

**Fig 5:**
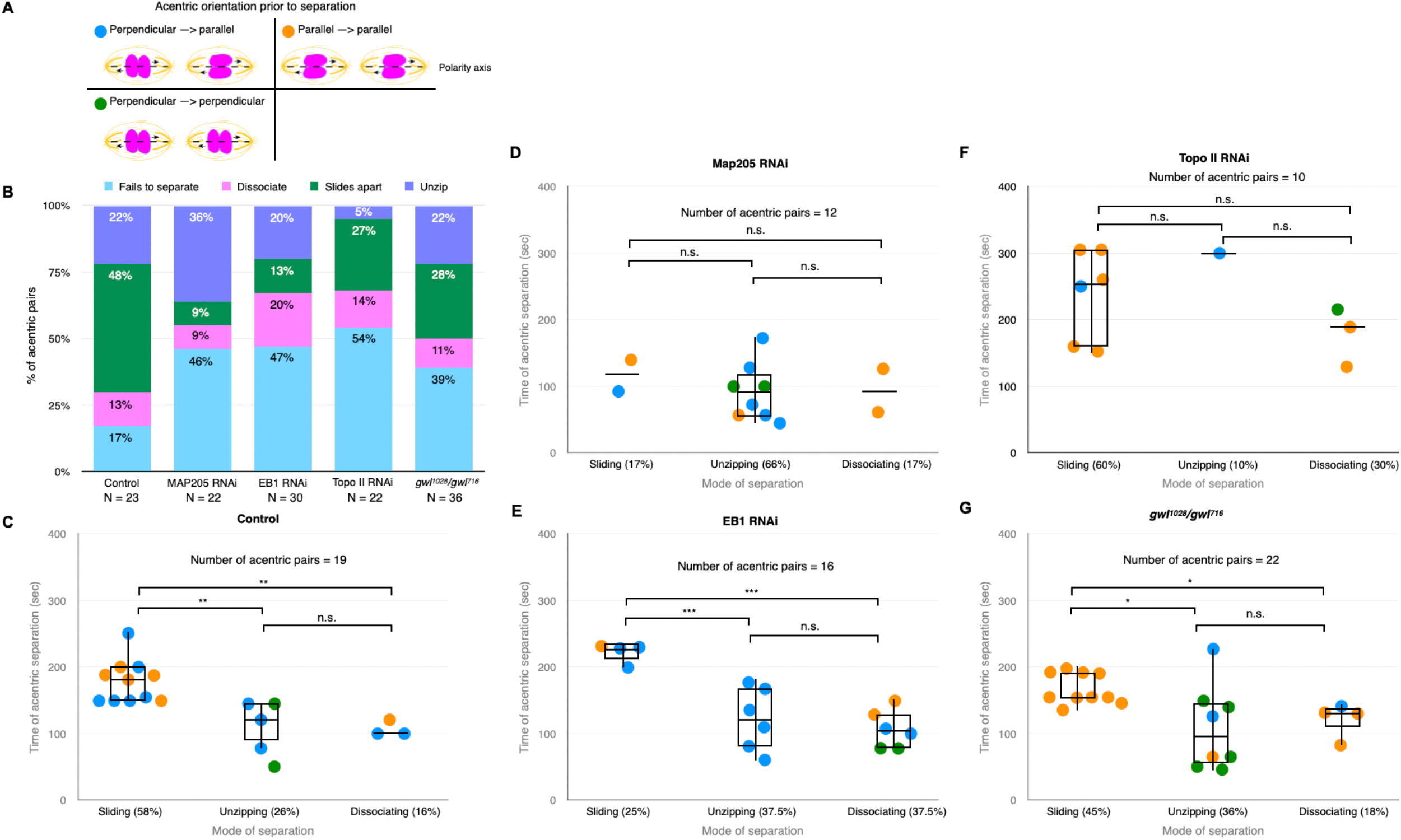
Knockdowns of EB1 and Map205 preferentially disrupt the sliding mode of acentric separation. (A) Model of different acentric orientations prior to separation. Polarity axis indicated by the dotted line. (B) Bar graph showing the percentage of acentric pairs that separate via simultaneous dissociation, sliding, or unzipping and the percentage of those that fail to separate. (C-G) Measurements of the frequency of each mode of acentric sister separation (x-axis) and the timing of acentric sister separation after the separation of intact chromosomes (y-axis). Each dot represents one acentric pair. Blue dots represent acentric pairs that were oriented perpendicular then parallel to the polarity axis prior to separation. Orange dots represent acentric pairs that were oriented parallel to the polarity axis prior to separation. Green dots represent acentric pairs that were oriented perpendicular to the polarity axis prior to separation. Boxes show interquartile ranges and lines show medians of the measured data. Asterisks indicate statistical significance (*P = 0.01, **P = 0.005, ***P = 0.009) when comparing the timing of acentric separation between separation modes. Statistical analysis was done using two-sided Mann-Whitney tests. Non-significant values had a P-value greater than 0.05.

**Table 4:** Average velocities of acentric segregation. Depicted below are the velocities of individual acentrics as they segregate during anaphase. Velocity is measured in nanometers per second. Segregation velocity is defined as beginning when an acentric is clearly observed as separated from its sister and ends when the acentric reaches the daughter nucleus. Asterisks indicate statistical significance (*P=0.006) as determined by a two-sided Mann-Whitney test.

| | Avg velocity ± SD<br>(nm/s) | N<br>(individual<br>acentrics) |
| --- | --- | --- |
| Control | 10 ± 4 | 26 |
| MAP205 RNAi | 12 ± 5 | 30 |
| EB1 RNAi | 10 ± 4 | 27 |
| Topo II RNAi | 12 ± 3 | 26 |
| *gwl<sup>1028</sup>/gwl<sup>716</sup>* | 8 ± 4 * | 32 |

Live analysis of acentric behavior in neuroblasts expressing EB1 RNAi revealed that 47% (14/30) of acentric sister pairs fail to separate (**Figure 4C, Figure 5E, Table 3**). The failure of separation results in acentric sisters segregating together to a single cell pole (**Figure 4B, Movie S3**). Additionally, the delay in acentric separation and their rate of poleward segregation were normal (**Figure 5E, Table 2, Table 4**). There was also not an increase in micronuclei formation (15% compared to 24% in controls) (**Figure 4D**). This indicates that the Map205 and EB1 knockdowns are specifically disrupting sister separation of acentrics and have no effect on the latter stages of acentric transmission.

As described earlier, cohesin removal on the intact chromosomes and acentrics occurs simultaneously, yet acentric sisters remain paired. This raised the possibility of a role for DNA catenation in maintaining acentric pairing. Previous work demonstrated that DNA catenations preserve sister chromatid cohesion in intact chromosomes until resolution by Topoisomerase II (Topo II) at the metaphase-anaphase transition [52]. Topo II decatenates intertwined DNA allowing sister chromatids to segregate to opposing poles of the cell. To test the role of DNA catenation in maintaining acentric sister pairing, we knocked down levels of Topo II specifically in the *Drosophila* neuroblast using the Gal4/UAS RNAi technique described above. Topo II knockdowns revealed that 54% (12/22) of sister acentrics fail to separate and subsequently segregate together to one pole (**Figure 6, Table 3, Movie S4**). Due to the lasting catenations between acentric sisters, the initiation of acentric separation was further delayed in neuroblasts expressing Topo II RNAi (Mann-Whitney test; P=0.003) (**Table 2**). Although acentric separation is disrupted, their poleward segregation rate and their rate of micronuclei formation were normal (**Figure 5F, Table 4, Figure 4D**).

**Fig 6:**
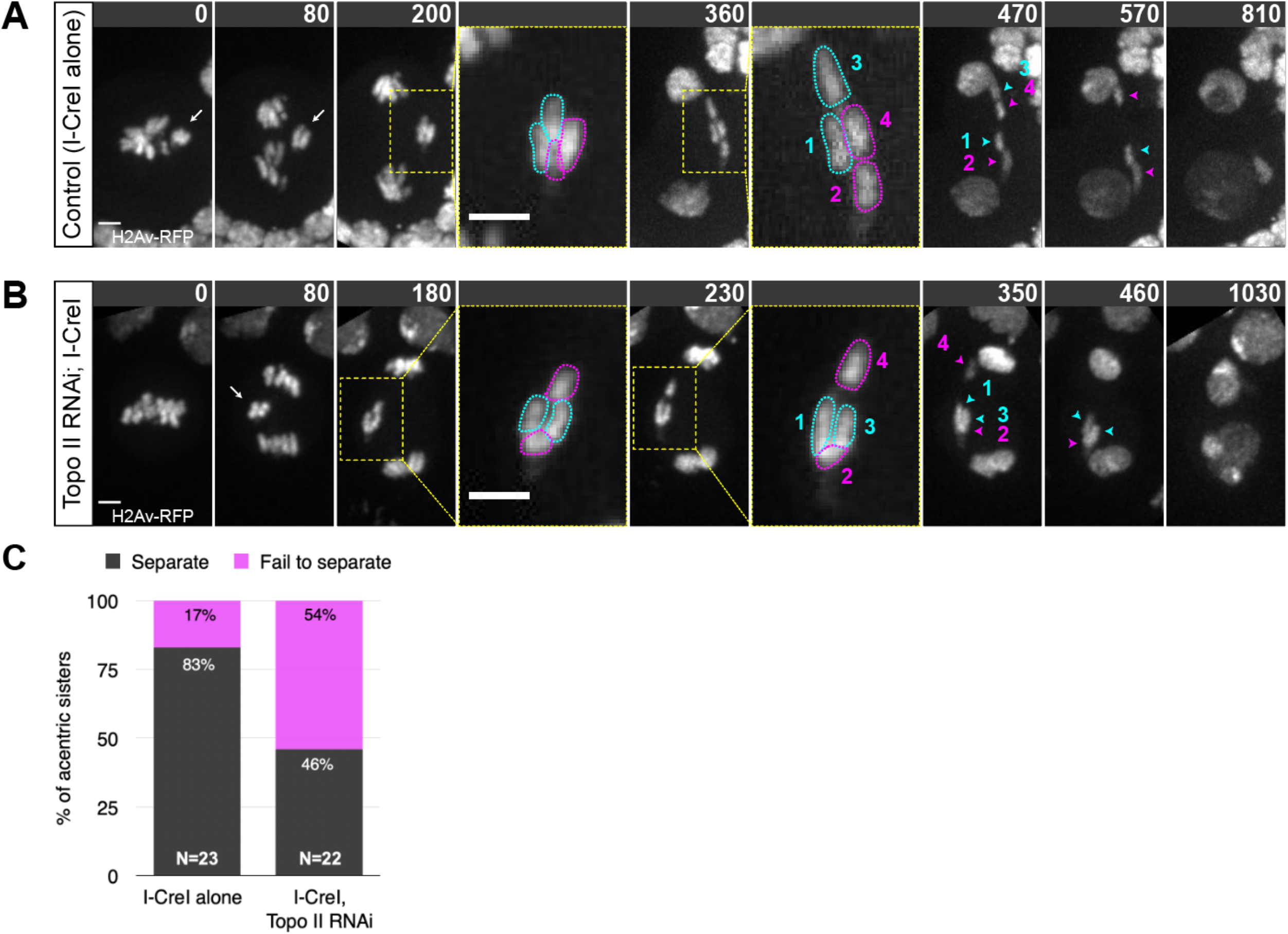
Acentrics appear to be held together by DNA catenations. (A) Still images of a time-lapse movie of a control neuroblast with I-CreI induced acentrics showing sister acentrics associated in early anaphase (white arrows), lagging behind and ultimately separating and segregating to opposite poles (cyan and magenta arrowheads). (B) Still images of a time-lapse movie of a neuroblast with partial knockdown of Topoisomerase II using RNAi and I-CreI induced acentrics. Acentrics associate and lag behind in early anaphase (white arrow), then fail to completely separate and segregate unequally (cyan and magenta arrowheads). Bars, 2 μm. Time in seconds. (Movie S4) (C) Bar graph showing the quantification of acentric behavior in control and Topo II RNAi neuroblasts with the failure of acentric sister separation in magenta and the successful, even separation of acentric sisters in gray.

The screen also yielded the PP2A inhibitor Greatwall kinase (*gwl*) that controls the timing and events of mitotic exit including sister chromatid separation [50]. Thus, we were particularly interested in the dynamics of acentric separation and segregation when the levels of *gwl* were reduced. This analysis uncovered that with reduced levels of *gwl*, 39% (14/36) of acentric sister pairs failed to separate from one another (**Figure 4C, Figure 5B, Table 3**). Additionally, the delay in acentric separation and the rate of micronuclei formation were normal (**Figure 5G, Table 2, Figure 4D**). However, the average velocity of acentrics while segregating during anaphase was significantly slower in *gwl* mutant background compared to acentrics in wildtype background (P=0.006; Mann-Whitney test) (**Table 4**).

### EB1, Map205, Greatwall, and Topo II differentially influence the dynamics of acentric separation

As described above, in a wild-type background acentrics exhibit three distinct patterns of separation: sliding, unzipping, and simultaneous dissociation. Sliding is the most frequent occurring 49% of the time, with unzipping and simultaneous dissociation occurring at frequencies of 31% and 20%, respectively (**Figure 3E**). To determine the role of the genes identified above in these distinct forms of acentric separation, we monitored the frequency of segregation patterns in RNAi knockdowns of the candidate genes. Of those sister acentrics that successfully separated, knockdowns of Map205, EB1, and Greatwall resulted in a decreased frequency of separation via sliding (17%, 25%, 45%, respectively) compared to the control (58%) (**Figure 5**). Additionally, Topoisomerase II resulted in a decreased the frequency of acentric separation via unzipping (10%) compared to the control (26%) (**Figure 5**).

To determine if microtubule plus-ends preferentially accumulate around acentrics, I-CreI-expressing neuroblasts were live imaged with EGFP-tagged EB1. Fluorescence intensities of both GFP and RFP were measured and corrected for brightness in control neuroblasts (N=5) and I-CreI-expressing neuroblasts (N=5). During mitosis, EB1 localizes along microtubules moving towards the plus-ends (**Figure 7 and Movie S5**). Previous studies have shown that EB1 is essential for generating antipolar forces on chromosomes [53, 54]. That finding together with our finding that EB1 is required for sister acentric separation motivated us to examine EB1 localization on the acentrics (**Figure 7**). We find the concentration of EB1 is not significantly increased on or near the acentrics (P > 0.05, two sample t-test; **Figure 7**). Thus, while EB1 and the acentrics co-localize, the pattern of EB1 comets is not altered in the presence of acentrics. While these data are consistent with a role for EB1 in acentric sister separation, there is not an increase in recruitment and accumulation of EB1 at the acentric.

**Fig 7:**
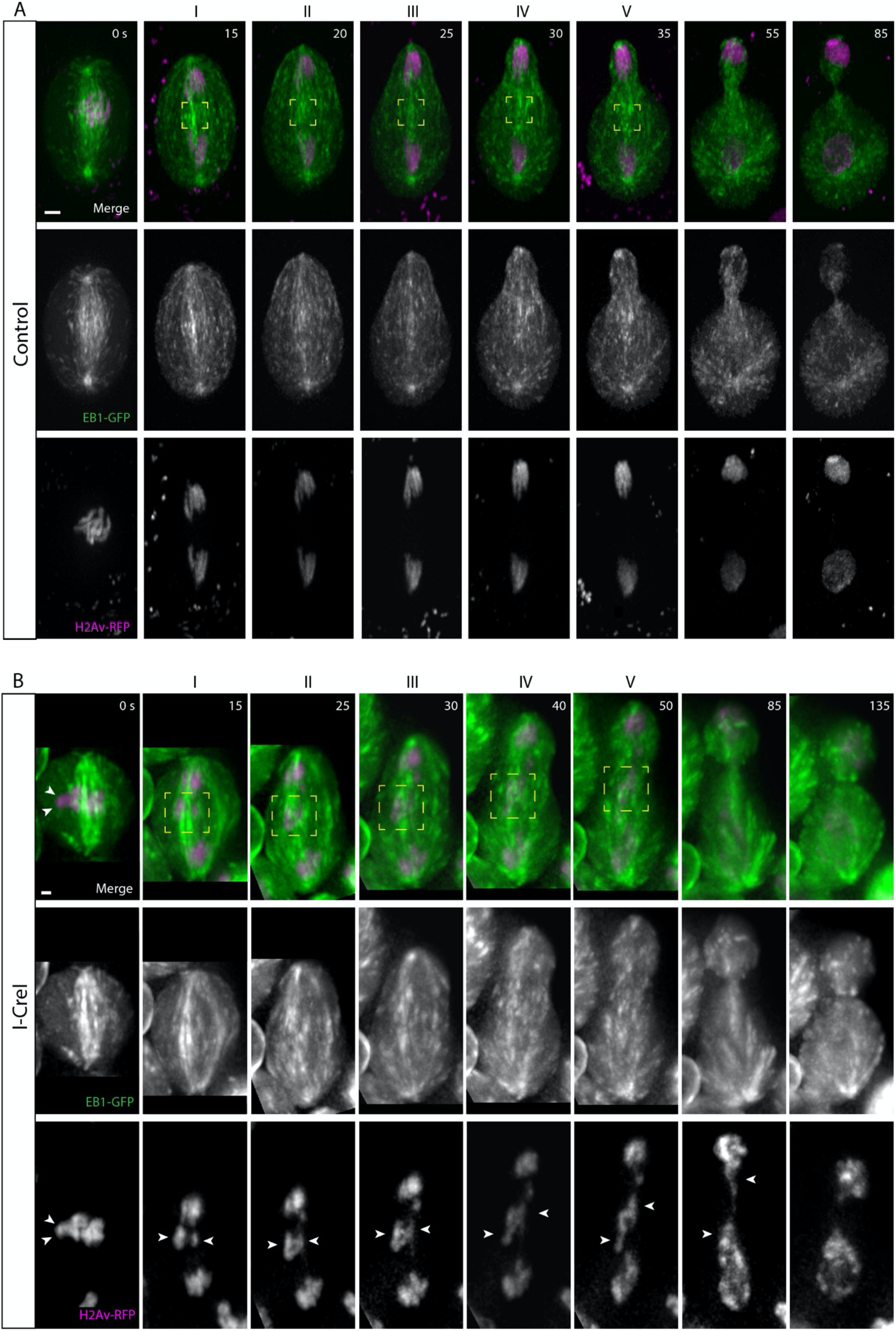
Acentrics travel poleward in late anaphase while associated with EB1. EB1 is in green and chromosomes are in magenta. (A) Still images from a time-lapse movie of a control neuroblast from metaphase (0 s) through telophase (85 s). (B) Still images from a time-lapse movie of a mitotic neuroblast with I-CreI induced acentrics from metaphase (0 s) through telophase (135 s) (Movie S5). The intact chromosomes and acentrics initiate sister separation at the 15 s and 40 s timepoints, respectively. Sister acentrics (white arrowheads) separate and move toward opposite cell poles while associated with EB1. (C) Line graph from a compilation of five control videos showing the corrected fluorescence intensities in arbitrary units (AU) of EB1 (green) and chromosomes (magenta). Corrected fluorescence intensities were calculated within the yellow boxes at the time points 15, 20, 25, 30, 35 seconds after anaphase onset. (D) Line graph from a compilation of five videos of I-CreI-expressing neuroblasts showing the corrected fluorescence intensities in arbitrary units (AU) of EB1 (green) and chromosomes (magenta). Corrected fluorescence intensities were calculated within the yellow boxes at the time points 15, 25, 30, 40, 50 seconds after anaphase. Bars, 2 μm. Time in seconds. Error bars represent SDs of the fluorescent intensities at all points tested.

## DISCUSSION

These studies are based on the unexpected finding that sister chromosome fragments lacking a kinetochore undergo relatively normal separation. We sought to identify the kinetochore independent forces driving acentric sister chromatid separation. A key finding is that while cohesin removal occurs simultaneously on intact and acentric chromosomes, sister separation of the latter is significantly delayed. An explanation for this delay comes from studies of intact chromosomes demonstrating that once cohesin is removed, sister acentrics remain held together through DNA catenations [52, 55, 56]. Catenations are concentrated at the centromeric DNA and opposing kinetochore microtubule interactions likely provide the resolving force [57–61]. Support for this conclusion comes from studies of chromosome rearrangements in which centromeric heterochromatin is displaced from the centromere [31]. Sister chromatin separation is specifically delayed in these regions resulting in localized stretching during anaphase. A likely consequence of being displaced from the centromere is that the ectopic heterochromatic regions no longer experience sufficient kinetochore forces required to efficiently resolve sister DNA catenations. In light of these studies, it is likely that acentric sisters remain associated well after separation of the intact chromosomes through DNA catenations. This conclusion is supported by our finding that reductions in the Topoisomerase II levels specifically disrupt acentric sister separation. As described below, our studies suggest both plus-end and lateral microtubule interactions with the acentrics provide the alternative force driving sister separation.

Intact chromosomes align perpendicular to the spindle. In contrast, we find acentric chromosomes align either perpendicular or parallel to the spindle. When aligned parallel to the spindle, acentrics travel with one tip leading towards the cell pole, possibly due to microtubule lateral interactions with acentrics. Without a kinetochore, a combination of lateral and plus-end microtubule interactions likely determines the final orientation of acentrics on the metaphase plate. In addition to the multiple orientations, acentrics undergo distinct patterns of sister separation that we have termed sliding, unzipping and simultaneous dissociation. Sliding of sisters past one another toward opposite poles is the most common mode of acentric separation. This mode occurs primarily when sister acentrics are oriented parallel to the spindle just prior to separation suggesting lateral microtubule interactions provide the force driving this mode of acentric separation (**Figure 3A, Figure 5C**). While sister separation of all acentrics is delayed relative to intact chromosomes, the delay is much more pronounced for acentrics that undergo separation by sliding. The delay may in part be due to the additional time it takes to establish the multiple lateral interactions required to generate sufficient separation force. It is likely chromokinesins provide the force driving separation, but these have yet to be identified.

In contrast to sliding, sister separation by unzipping occurs primarily when the acentrics are oriented perpendicular to the spindle just prior to separation suggesting a limited role for lateral microtubule interactions (**Figure 3B, Figure 5C**). During unzipping, acentrics generally initiate separation at their broken ends with completion of separation occurring at the telomeres. The delay of acentric separation by unzipping is much less than the delay observed for separation by sliding. Previous studies demonstrated that the acentric and its centric partner are connected via a DNA tether [12]. The preference for initiation of separation at the broken-ends suggests that a DNA tether connecting the broken end to the kinetochore-bearing fragment may provide the initial separation force (**Figure 3B**). While this tether is not thought to provide the force driving acentric segregation [15], because of its association with the broken end of the acentric, it may drive the initial stage of unzipping. In contrast to separation by sliding, telomeres are always aligned in sisters that separate by unzipping (**Figure 3E, Figure 3F**). This suggests maintaining gene-for-gene sister pairing is essential for this form of separation.

The third mode of acentric separation, simultaneous dissociation, is characterized by a synchronized separation along the entire length of sister acentrics (**Figure 3C**). While sister separation via simultaneous dissociation favors the perpendicular orientation, this bias is much less dramatic compared to the unzipping mode (**Figure 5C**). The most distinguishing feature of this form of separation is the high frequency in which sister acentrics aligned with their telomeres opposed (33% compared to 14% and 0% for sliding and unzipping separation, respectively) (**Figure 3F**). This high frequency of unaligned telomeres suggests weak connections between sisters and that sister pairing is not required.

Synthetic lethal screens have proven an effective means of identifying factors required for the successful mitotic transmission of acentric chromosome fragments [12, 15]. These screens have identified proteins required for poleward transmission and final incorporation of acentrics into the telophase nucleus. Here we focused on identifying genes required for successful separation of sister acentrics. Of the 117 candidate genes screened, we identified 23 RNAi knockdowns/mutations that resulted in a significant lethality upon acentric induction (**Table 1**). Live analysis secondary screening revealed knockdowns of EB1 and Map205, and the cell cycle kinase Greatwall resulted in dramatic defects in acentric sister separation. Live imaging also revealed reducing levels of Topoisomerase II severely disrupted sister acentric separation.

Given that previous studies demonstrated the chromokinesin Klp3A is required for acentric poleward transport [15], it is not surprising that microtubule-associated proteins also play a key role in separation of acentric sisters. Reduced levels of EB1 and Map205 greatly increase the frequency of failed sister separation. EB1 is a plus-end tracking protein that associates with a number of regulatory proteins and is essential for generating anti-polar forces on the chromosomes [62]. EB1 knockdowns most dramatically reduce the frequency of acentric separation via sliding. In addition, for those acentric sisters that did separate by sliding in EB1 knockdowns, separation was greatly delayed. As lateral interactions between the microtubules and chromatin are likely to drive sister separation by sliding, we suspect plus-end directed EB1 forces may play a role in orienting the acentric in order to establish lateral interactions. Support for this idea comes from the finding of a synthetic lethal interaction between I-CreI induction and reductions in the levels of Nod, a non-motile chromokinesin that associates with EB1 and is involved in chromosome segregation [15, 63–65]. Surprisingly, a previous study in our lab found no effect on acentric segregation in mutations in Nod [15]. Examining movies from this analysis revealed no significant difference in acentric sister separation in a loss-of-function *nod* mutant compared to the control (**Table S2**). Map205 is a microtubule-associated protein required for targeting Polo kinase to spindle microtubules [42]. Thus, it is likely that the effect of the Map205 knockdowns on acentric sister separation is through disruption of Polo localization. Previous studies demonstrated that Polo localizes to the DNA tether associated with the acentric and Polo knockdowns disrupted acentric transmission with sister acentrics remaining on the metaphase plate [12]. As with EB1, reductions in Map205 levels dramatically reduce the frequency of acentric separation by sliding suggesting a disruption in lateral interactions between the acentrics and the microtubules. While much evidence indicates that Polo functions at the centromere during sister separation, our studies of the effects of Map205 and Polo knockdowns suggest that it is also involved in promoting chromatid/spindle interactions that are independent of the centromere. Reducing the levels of Topoisomerase II leads to a significant increase in failed separation of sister acentrics, supporting the conclusion that once cohesin is removed sister acentrics remain held together through DNA catenation. We suspect that reduced Topoisomerase II levels disrupt sister separation of the acentrics more profoundly than that of intact chromosomes because the latter primarily rely on kinetochore forces to resolve DNA catenations. Of the four mutants analyzed through live analysis, reductions in Topoisomerase levels had the greatest effect of the unzipping mode of acentric sister separation. This result is interesting given our finding that the unzipping mode of acentric separation is likely the most reliant on sister pairing and consequently are likely to be highly catenated.

Taken together, these data support a model in which multiple forces drive the separation of sister acentrics. Acentrics that separate by sliding and unzipping tend to be oriented parallel and perpendicular to the spindle, respectively. Unzipping could be initially driven by the DNA tether, which connects the acentric to its centric partner, to initially separate one end of the acentrics. Then, a combination of Topoisomerase II activity and microtubule forces could facilitate the separation of the other end of the acentrics. Because the partial knockdown of Topoisomerase II leads to a decreased frequency of acentrics unzipping, we hypothesize that the resolution of DNA catenations by Topoisomerase II underlies the unzipping mode of separation. Due to the finding that the tether does not appear to provide segregation forces on the acentrics [15], sliding is likely driven by lateral microtubule interactions and possibly motor proteins. This is further supported by the orientation of acentrics parallel to the axis of polarity and the association of EB1 and bundling of microtubules around acentrics in early anaphase. It remains unclear if acentrics are directly interacting with microtubules or if microtubule-associated proteins are mediating the interaction with acentrics. There is not an obvious mechanism for the simultaneous dissociation of sister acentrics and is likely driven by a combination of factors (**Figure 8**). EB1 and Map205 may be required to establish microtubule interactions with the acentric, while in the absence of a kinetochore high levels of Topoisomerase are needed to resolve DNA catenations between sisters.

**Fig 8:**
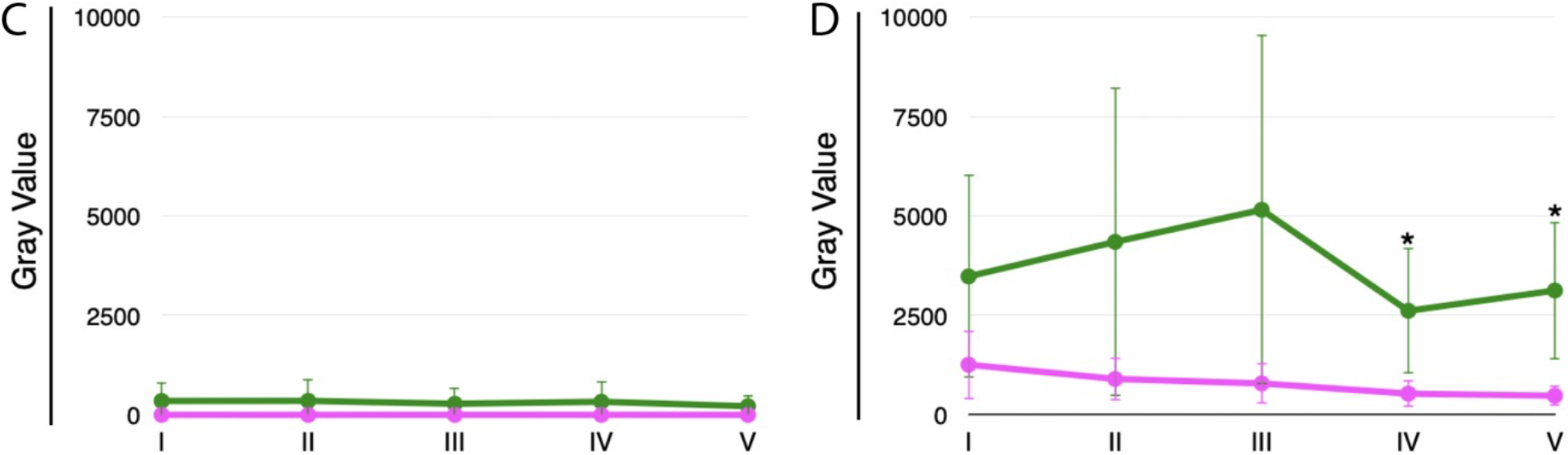
Schematic showing distinct modes of acentric separation. At some point during the metaphase-to-anaphase transition, acentrics that separate by sliding orient parallel to the spindle and slide past one another. This is likely driven by lateral associations between the microtubules and acentrics. The unzipping mode occurs when separation initiates at the broken end followed by separation of sister telomeres. It may be that the initial separation is driven by the DNA tether connecting the centric and acentric fragments. Separation across the entire length of the acentric is referred to as simultaneous dissociation. It may be that the DNA tether and acentric-microtubule interactions contribute equally for sister acentrics that separate by simultaneous dissociation. (Chromosomes in magenta, microtubules in green, cohesin in cyan, telomeres in orange).

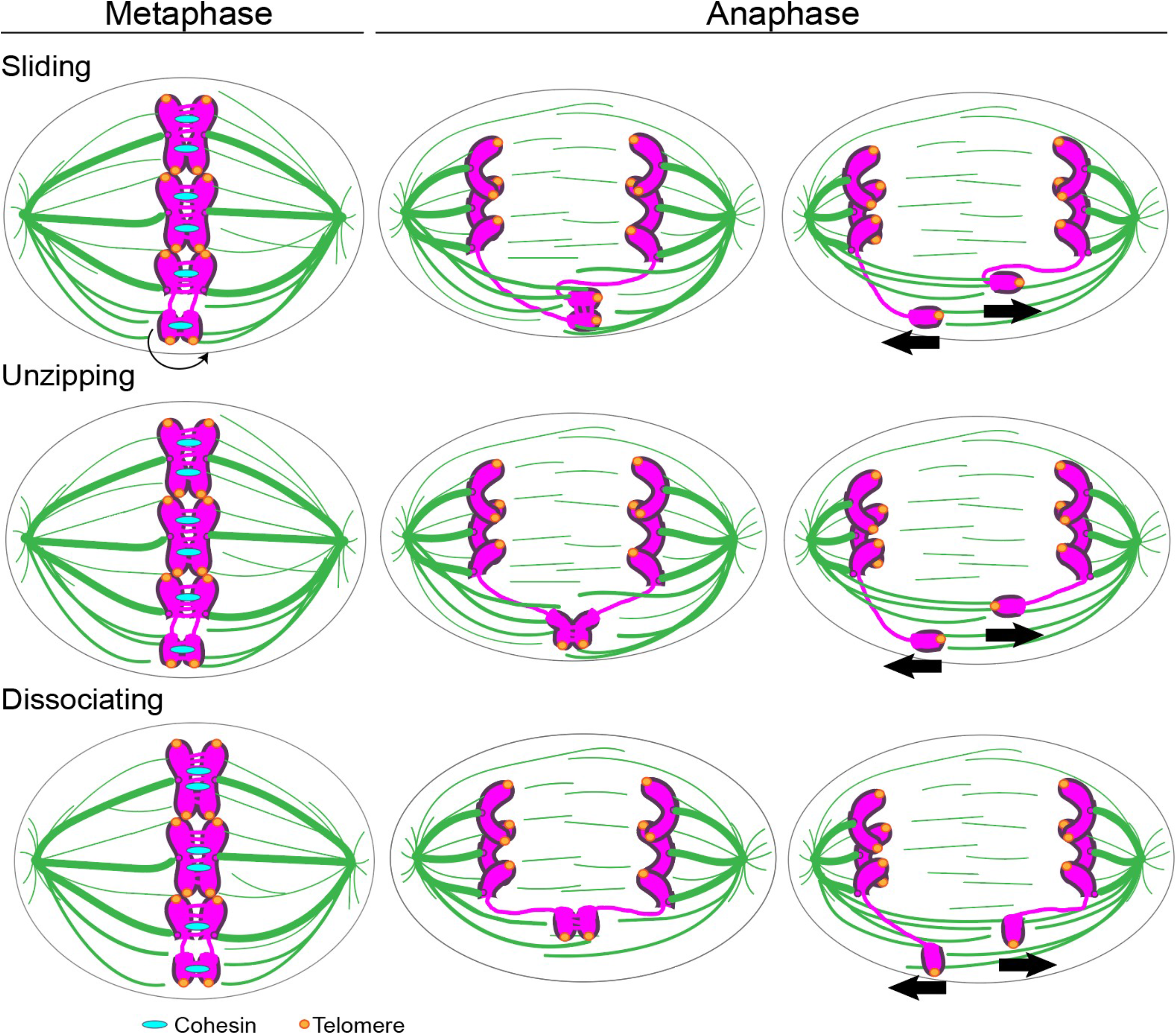

It should be noted that although these mutants resulted in a high frequency of failed acentric sister separation, we did not observe an equivalent increase in micronuclei. In contrast, mutants that disrupted acentric poleward transport or the final stages of incorporation into daughter nuclei resulted in an accompanying increase in micronuclei [66–68]. This indicates that the knockdowns in the genes identified in this study specifically disrupt acentric sister separation as the subsequent transmission occurs normally to allow incorporation of the acentric into the daughter nuclei. This interpretation is in accord with a model that acentrics experience different forces at different stages of their separation, transmission, and incorporation into daughter nuclei [68].

There are numerous examples of lateral interactions between spindle microtubules and intact chromosomes indicating that non-kinetochore forces influence anaphase chromosome kinetics [69–71]. However, it has been difficult to pursue the underlying mechanisms because of the dominance of kinetochore forces during anaphase. In analogy with studies of spindle formation in cells lacking centrosomes that led to the discovery of unsuspected, yet conserved, chromosome-based mechanisms of spindle assembly, examining the mitotic behavior of acentric chromosomes without a kinetochore will provide insights into the kinetochore independent forces acting on intact chromosomes [70, 71].

## MATERIALS AND METHODS

### Fly Stocks

All stocks were raised on standard *Drosophila* media at room temperature (20-22°C) as previously described [72]. For generating acentrics, a transgenic fly line bearing the I-CreI endonuclease under heat-shock 70 promoter were kindly provided by Kent Golic at The University of Utah. For synthetic lethality screen, the ubiquitous Gal4 driver under the control of an actin enhancer (Act5) was used (#25708 from Bloomington). Dominant negative allele of ISWI was kindly provided by John Tamkun at UC Santa Cruz. Greatwall hypomorphs (*gwl*^*1080, 716, 180, and 2790*^) were kindly provided by Michael Goldberg at Cornell University. The line with *rad21-EGFP* transgene were kindly provided by Stefan Heidmann at University of Bayreuth.

### Synthetic Lethality Screen

Third instar progeny with genotype Act5-Gal4/+; I-CreI, Sb/UAS-RNAi were collected from parental genotypes Act5-Gal4/CyO-GFP; I-CreI, Sb/TM6B and were heat shocked for 1.5 hours at 37°C (unless otherwise indicated). After heat shock, the vials were set-aside at room temperature for 10-15 days until adult flies emerged. Synthetic lethality was calculated as the % of larvae that develop into adulthood [12, 15]. Control progeny with genotype Act5-Gal4/+; Sb/UAS-RNAi from parents with the genotype Act5-Gal4/CyO-GFP; Sb/TM6B and UAS-RNAi were heat-shocked for 1.5 hours at 37°C.

### Live analysis of acentric behavior in *Drosophila* third instar neuroblasts

As previously described, acentric chromosome fragments were induced by I-CreI expression (under heat shock 70 promoter) in 3^rd^ instar larvae by a 1-hour 37°C heat shock followed by a 1-hour recovery period at room temperature [12]. The larval brains from third instar larvae were dissected in PBS and then transferred to a slide with 20 μl of PBS. A coverslip was dropped on PBS with brain and the excess PBS was wicked out from edge of coverslip to induce squashing of brain between slide and coverslip. For live analysis, the edge of coverslip was sealed with halocarbon and was imaged as described below. Neuroblast divisions in all images were from female 3^rd^ instar larvae.

### Microscopy and image acquisition

#### Wide-field microscopy

Time-lapse imaging for Figure 4C, 4D, 5C, and 6 were performed using a Leica DM16000B wide-field inverted microscope equipped with a Hamamatsu electron-multiplying charge coupled device camera (ORCA 9100-02) with a binning of 1 and a 100x Plan-Apochromat objective with NA 1.4. Successive time points were filmed at 20 s. RFP (585 nm) and GFP (508 nm) fluorophores were imaged. Samples were imaged in PBS and at room temperature (20-22°C). Widefield images were acquired with Leica Application Suite Advanced Fluorescence Software and 3D deconvolved using AutoQuant X2.2.0 software.

#### Spinning-disk microscopy

Images in Figures 1, 2, 3, 4, 6, and 7 were acquired with an inverted Nikon Eclipse TE2000-E spinning disk (CSLI-X1) confocal microscope equipped with a Hamamatsu electron-multiplying charge coupled device camera (ImageEM X2) with a 100X 1.4 NA oil-immersion objective. Samples were imaged in PBS and at room temperature (20-22°C). Images were acquired with MicroManager 1.4 software. Time-lapse fluorescent images of neuroblasts divisions were done with 120 and 100 ms exposures for GFP and RFP respectively with 0.5 μm Z-steps. Time-lapse videos with both GFP and RFP were done every 5 to 9 seconds and time-lapse movies with RFP alone were done every 5 seconds. Figures were assembled in Adobe Illustrator. Selected stills (both experimental and control) were processed with ImageJ (http://rsb.info.nih.gov/ij/).

### Measurements

In Figure 2C, relative fluorescence intensities of chromosomes (H2Av-RFP) and cohesin (Rad21-EGFP) were done using the plot profile function in ImageJ of the region outlined around acentrics and the main mass of chromosomes. In Figures 7C and 7D, relative fluorescence intensities of acentrics (H2Av-RFP) and EB1 (EB1-EGFP) were done using the plot profile function in ImageJ of the region outlined around acentrics in I-CreI expressing neuroblasts and the region outlined around the spindle midzone in control neuroblasts. For statistical analyses, unpaired two-sided t-tests and two-sided Mann-Whitney-Wilcoxon tests were used. Unpaired two-sided t-tests were performed in Prism Version 8 (GraphPad Software). Two-sided Mann-Whitney-Wilcoxon tests were performed in R (R Core Team) and Prism Version 8 (GraphPad Software).

## Supporting information

Supplemental Movies Description

## ACKNOWLEDGEMENTS

We would like to thank Dr. Susan Strome, Dr. William Saxton, and Dr. John Tamkun for use of their equipment and providing reagents. We thank Dr. Benjamin Abrams for his advice and technical expertise regarding microscopy experiments. We thank Dr. Kent Golic for providing us with the transgenic fly line bearing the I-CreI endonuclease. We thank Dr. Michael Goldberg for providing us with Greatwall hypomorph flies. We thank Dr. Stefan Heidmann for providing us with the transgenic fly line bearing *rad21-EGFP*. We thank H. Vicars’ service dog Teddy Bear for his guidance and support.

Funding for these studies was provided by National Institutes of Health grant NIH1RO1GM120321 awarded to W. Sullivan.

The authors declare no competing financial interests.

## Notes

### Competing Interest Statement

The authors have declared no competing interest.

### Summary of Updates

Figures, statistical analysis, supplemental files, and manuscript text have been updated

