## Supplemental Tables for "Kinetochore-independent mechanisms of sister chromosome separation": Supplemental Table 1 and 2.pdf

### Supplemental Table: Synthetic Lethality Screen

| Gene (RNAi) | Stock # | I-CreI | Number of experiments | Number of Larvae per experiment | Heat Shock (1.5 hours) | Normal Adults per experiment | Mutant Phenotype per experiment | % Survival (Avg ± St Dev) | Mutant Wing Phenotype (%) |
|---|---|---|---|---|---|---|---|---|---|
| WT (control) | . | Yes | 9 | 28,41,32,10,25,25, 7,13,13 | Yes | 24,41,11,9,21,15, 7,9,12 | | 80 ± 21 | 0 |
| WT (control) | . | No | 9 | 13,14,33,20,26,25, 8,29,26 | No | 13,13,26,17,25,24,7,26, 25 | | 91 ± 7 | 0 |
| Acf1 | 35575 | Yes | 1 | 25 | Yes | 17 | 1 with wing spot | 68 | 4 |
| Acf1 | 35575 | No | 1 | 13 | No | 9 | | 69.23 | 0 |
| Ada2b | 35334 | Yes | 1 | 25 | Yes | 18 | | 72 | 0 |
| Ada2b | 35334 | No | 1 | 17 | No | 16 | | 94.12 | 0 |
| ald | 35283 | Yes | 1 | 25 | Yes | 13 | | 52 | 0 |
| ald | 35283 | No | 1 | 25 | No | 22 | | 88 | 0 |
| ald | 36658 | Yes | 1 | 21 | Yes | 14 | | 66.67 | 0 |
| ald | 36658 | No | 1 | 12 | No | 9 | | 75 | 0 |
| Ald | 26301 | Yes | 1 | 25 | Yes | 0 | Lethal after 3rd Instar | . | . |
| Ald | 26301 | No | 1 | 25 | No | 0 | Lethal after 3rd Instar | . | . |
| asf1 | 35273 | Yes | 1 | 25 | Yes | 20 | 2 wing notch | 80 | 8 |
| asf1 | 35273 | No | 1 | 25 | No | 20 | 1 deformed wing | 80 | 4 |
| asp | 28741 | Yes | 1 | 25 | Yes | 13 | | 52 | 0 |
| asp | 28741 | No | 1 | 15 | No | 15 | | 100 | 0 |
| asp | 35224 | Yes | 1 | 25 | Yes | 11 | 5 Wing Notch | 44 | 20 |
| asp | 35224 | No | 1 | 25 | No | 24 | 1 Deformed wing | 96 | 4 |
| barr | 34068 | Yes | 1 | 25 | Yes | 0 | Lethal after 3rd Instar | . | . |
| barr | 34068 | No | 1 | 25 | No | 0 | Lethal after 3rd Instar | . | . |
| brm | 34520 | Yes | 1 | 16 | Yes | 12 | | 75 | 0 |
| brm | 34520 | No | 1 | 21 | No | 21 | | 100 | 0 |
| brat | 34646 | . | 1 | 0 | . | | Lethal before 3rd instar | . | . |
| brat | 34646 | . | 1 | 0 | . | | Lethal before 3rd instar | . | . |
| Bub1 | 35260 | Yes | 1 | 25 | Yes | 22 | | 88 | 0 |
| Bub1 | 35260 | No | 1 | 20 | No | 19 | | 95 | 0 |
| BubR1 | 35329 | Yes | 6 | 25,24,36,33,25,25 | Yes | 1,0,1,0,0,0 | 1 wrinkled wing | 1 ± 2 | 4 |
| BubR1 | 35329 | No | 3 | 25,16,28 | No | 14,4,3 | | 31 ± 23 | 0 |
| Bub3 | 32989 | Yes | 1 | 25 | Yes | 0 | Adults dead inside pupae case | . | . |
| Bub3 | 32989 | No | 1 | 25 | No | 0 | Adults dead inside pupae case | . | . |
| Caf1 | 34069 | Yes | 1 | 26 | Yes | 0 | Lethal after 3rd Instar | . | . |
| Caf1 | 34069 | No | 1 | 23 | No | 0 | Lethal after 3rd Instar | . | . |
| caf1-180 | 32478 | Yes | 1 | 39 | Yes | 0 | Lethal after 3rd Instar | . | . |
| caf1-180 | 32478 | No | 1 | 25 | No | 0 | Lethal after 3rd Instar | . | . |
| Cap | 33431 | Yes | 1 | 25 | Yes | 0 | Lethal after 3rd Instar | . | . |
| Cap | 33431 | No | 1 | 13 | No | 0 | Lethal after 3rd Instar | . | . |
| CAP-D2 | 31478 | Yes | 1 | 25 | Yes | 0 | Lethal after 3rd Instar | . | . |
| CAP-D2 | 31478 | No | 1 | 32 | No | 0 | Lethal after 3rd Instar | . | . |
| CAP-D2 | 31326 | Yes | 1 | 17 | Yes | 1 | | 5.88 | 0 |
| CAP-D2 | 31326 | No | 1 | 12 | No | 6 | | 50 | 0 |
| car | 34007 | Yes | 1 | 25 | Yes | 19 | | 76 | 0 |
| car | 34007 | No | 1 | 23 | No | 21 | | 91.3 | 0 |
| Cdc2 | 36117 | Yes | 1 | 28 | | 0 | Lethal after 3rd Instar | . | . |
| Cdc2 | 36117 | No | 1 | 18 | | 0 | Lethal after 3rd Instar | . | . |
| Cenp-C | 34692 | Yes | 1 | 25 | Yes | 0 | Lethal after 3rd Instar | . | . |
| Cenp-C | 34692 | No | 1 | 22 | No | 0 | Lethal after 3rd Instar | . | . |
| Cenp-C | 34699 | Yes | 1 | 25 | Yes | 0 | Lethal after 3rd Instar | . | . |
| Cenp-C | 34699 | No | 1 | 25 | No | 0 | Lethal after 3rd Instar | . | . |
| Cenp-C | 26311 | Yes | 1 | 0 | Yes | 0 | Lethal before 3rd Instar | . | . |
| Cenp-C | 26311 | No | 1 | 0 | No | 0 | Lethal before 3rd Instar | . | . |
| Chb | 34669 | Yes | 1 | 27 | Yes | 0 | Lethal after 3rd Instar | . | . |
| Chb | 34669 | No | 1 | 22 | Yes | 0 | Lethal after 3rd Instar | . | . |
| Chd1 | 34665 | Yes | 3 | 20,24,25 | Yes | 0,7,0 | | 10 ± 17 | 0 |
| Chd1 | 34665 | No | 5 | 20,23,13,24,19 | Yes | 6,9,5,7,5 | | 33 ± 6 | 7 |
| chd1 | 35240 | Yes | 1 | 25 | Yes | 12 | 1 with wing notch | 48 | 4 |
| chd1 | 35240 | No | 1 | 23 | No | 22 | | 95.65 | 0 |
| Chrac-14 | 35652 | Yes | 1 | 25 | Yes | 15 | 2 with wing spots | 60 | 8 |
| Chrac-14 | 35652 | No | 1 | 16 | No | 16 | | 100 | 0 |
| Chrac-14 | 31052 | Yes | | 25 | Yes | 12 | | 48 | 0 |
| Chrac-14 | 31052 | No | | 25 | No | 25 | | 100 | 0 |
| Cmet | 35816 | Yes | 1 | 18 | Yes | 0 | Lethal after 3rd Instar | . | . |
| Cmet | 35816 | No | 1 | 25 | No | 0 | Lethal after 3rd Instar | . | . |
| Cp190 | 33903 | Yes | 1 | 17 | Yes | 2 | | 11.8 | 0 |
| Cp190 | 33903 | No | 1 | 12 | Yes | 3 | most dead in bottom food or "push-pop" phenotype | 25 | 0 |
| CtBP | 32889 | Yes | 1 | 25 | Yes | 0 | Lethal after 3rd Instar | . | . |
| CtBP | 32889 | No | 1 | 24 | No | 0 | Lethal after 3rd Instar | . | . |
| D1 | 33655 | Yes | 1 | 0 | Yes | 0 | Lethal before 3rd Instar | . | . |
| D1 | 33655 | No | | 0 | No | 0 | Lethal before 3rd Instar | . | . |
| Dhc | 36698 | Yes | 1 | 0 | . | . | Lethal before 3rd instar | . | . |
| Dhc | 36698 | No | 1 | 0 | . | . | Lethal before 3rd instar | . | . |
| Dhc | 28749 | Yes | 1 | 0 | . | . | Lethal before 3rd instar | . | . |
| Dhc | 28749 | No | 1 | 0 | . | . | Lethal before 3rd instar | . | . |
| dlg | 33620 | . | 1 | 0 | . | | Lethal before 3rd instar | . | . |
| dlg | 33620 | . | 1 | 0 | . | | Lethal before 3rd instar | . | . |
| dpn | 26320 | Yes | 1 | 25 | Yes | 17 | | 68 | 0 |
| dpn | 26320 | No | 1 | 24 | No | 22 | | 91.67 | 0 |
| EB1 | 28605 | Yes | 7 | 10,8,7,16,6,14,25 | Yes | 8,6,1,4,2,5,12 | 11 with wing notches | 44 ± 25 | 21 ± 6 |
| | | | | | | | 1 deformed wing | | |
| EB1 | 28605 | No | 4 | 19,4,5,10 | Yes | 18,4,5,8 | | 94 ± 9 | 0 |
| E(bx) | 33658 | Yes | 1 | 0 | . | . | Lethal before 3rd instar | . | . |
| E(bx) | 33658 | No | 1 | 0 | . | . | Lethal before 3rd instar | . | . |
| Futch | 40839 | Yes | 1 | 9 | Yes | 8 | 1 wing notch | 88.9 | 11 |
| Futch | 40839 | No | 1 | 7 | Yes | 7 | | 100 | 0 |

| Gene (RNAi) | Stock # | I-CreI | Number of experiments | Number of Larvae per experiment | Heat Shock (1.5 hours) | Normal Adults per experiment | Mutant Phenotype per experiment | % Survival (Avg ± St Dev) | Mutant Wing Phenotype (%) |
|---|---|---|---|---|---|---|---|---|---|
| **grp** | 27277 | Yes | 1 | 25 | Yes | 20 | 1-Notched wing, 1-Wing Spot, 2-Blistered wing | 80 | 16 |
| **grp** | 27277 | No | 1 | 15 | No | 15 | | 100 | 0 |
| **grp** | 36685 | Yes | 1 | 0 | Yes | 0 | Lethal before 3$^{rd}$ instar | . | . |
| **grp** | 36685 | No | 1 | 0 | Yes | 0 | Lethal before 3$^{rd}$ instar | . | . |
| **Grip75** | 31215 | Yes | 1 | 23 | Yes | 20 | 1 male notch | 87 | 4 |
| **Grip75** | 31215 | No | 1 | 7 | Yes | 7 | | 100 | 0 |
| **Grip84** | 33458 | Yes | 1 | 21 | Yes | 21 | | 100 | 0 |
| **Grip84** | 33458 | No | 1 | 15 | Yes | 14 | | 93.3 | 0 |
| **gwl** | 35212 | Yes | 1 | 25 | Yes | 22 | 1 Major wing spot | 88 | 4 |
| **gwl** | 35212 | No | 1 | 18 | No | 17 | | 94.44 | 0 |
| **gwl** | 34525 | Yes | 5 | 23,16,14,39,20 | Yes | 0,0,0,1,1 | | 2 ± 2 | 0 |
| **gwl** | 34525 | No | 5 | 21,22,22,21,13 | Yes | 9,5,4,5,3 | 44 wing notches | 26 ± 10 | 44 ± 19 |
| **hairy** | 34326 | Yes | 1 | 25 | Yes | 19 | | 76 | 0 |
| **hairy** | 34326 | No | 1 | 25 | No | 25 | | 100 | 0 |
| **Hira** | 35346 | Yes | 1 | 25 | Yes | 18 | | 72 | . |
| **Hira** | 35346 | No | 1 | 26 | No | 26 | | 100 | . |
| **His2Av** | 34844 | Yes | 1 | 0 | Yes | 0 | Lethal prior to 3rd Instar | . | 0 |
| **His2Av** | 34844 | No | 1 | 0 | No | 0 | Lethal prior to 3rd Instar | . | 0 |
| **His3.3B** | 34940 | Yes | 1 | 26 | Yes | 24 | 1 with blistered wing | 92.31 | 4 |
| **His3.3B** | 34940 | No | 1 | 25 | No | 21 | | 84 | 0 |
| **HmgZ** | 26219 | Yes | 1 | 22 | Yes | 17 | 2 notched wing | 77.27 | 9 |
| **HmgZ** | 26219 | No | 1 | 25 | No | 16 | | 64 | 0 |
| **HmgD** | 31344 | Yes | 1 | 25 | Yes | 25 | | 100 | 0 |
| **HmgD** | 31344 | No | 1 | 25 | No | 23 | | 92 | 0 |
| **HP1b** | 32401 | Yes | 1 | 20 | Yes | 13 | | 65 | 0 |
| **HP1b** | 32401 | No | 1 | 25 | No | 21 | | 84 | 0 |
| **HP1C** | 33962 | Yes | 1 | 25 | Yes | 15 | 2 with wing Notch | 60 | 8 |
| **HP1C** | 33962 | No | 1 | 25 | No | 22 | | 88 | 0 |
| **HP1e** | 34863 | Yes | 1 | 20 | Yes | 13 | | 65 | 0 |
| **HP1e** | 34863 | No | 1 | 25 | No | 25 | | 100 | 0 |
| **hyx** | 31722 | . | | 25 | . | 0 | Lethal prior 3$^{rd}$ instar | . | . |
| **hyx** | 31722 | . | | 25 | . | 0 | Lethal prior 3$^{rd}$ instar | . | . |
| **hyx** | 35238 | Yes | 1 | 25 | Yes | 2 | 1 deformed wing | 8 | 4 |
| **hyx** | 35238 | No | 1 | 25 | No | 22 | 1 deformed wing | 88 | 4 |
| **ial** | 28691 | Yes | 1 | 25 | Yes | 0 | Lethal after 3$^{rd}$ Instar | . | . |
| **ial** | 28691 | No | 1 | 13 | No | 0 | Lethal after 3$^{rd}$ Instar | . | . |
| **Incenp** | 35366 | Yes | 1 | 20 | Yes | 0 | Lethal after 3$^{rd}$ Instar | . | . |
| **Incenp** | 35366 | No | 1 | 17 | No | 0 | Lethal after 3$^{rd}$ Instar | . | . |
| **Ino80** | 37473 | Yes | 1 | 15 | Yes | 14 | | 93.33 | 0 |
| **Ino80** | 37473 | No | 1 | 10 | No | 10 | | 100 | 0 |
| **Ino80** | 33708 | Yes | 1 | 26 | Yes | 26 | | 100 | 0 |
| **Ino80** | 33708 | No | 1 | 14 | No | 14 | | 100 | 0 |
| **Iswi** | 32845 | Yes | 1 | 25 | Yes | 1 | 1 wing spot | 4 | 4 |
| **Iswi** | 32845 | No | 1 | 6 | Yes | 0 | | 0 | 0 |
| **Iswi** | 31111 | Yes | 1 | 40,27,36,25,25,25 | Yes | 17,8,26,4,11,14 | 1 wing notch, 3 wing spot | 43 ± 20 | 16 |
| **Iswi** | 31111 | No | 3 | 42,7,40 | Yes | 20,6,29 | | 69 ± 19 | 0 |
| **Klp68D** | 29410 | Yes | 1 | 24 | Yes | 7 | | 29.17 | 0 |
| **Klp68D** | 29410 | No | 1 | 21 | No | 12 | | 57.14 | 0 |
| **Klp67A** | 35606 | Yes | 1 | 19 | Yes | 10 | | 52.63 | 0 |
| **Klp67A** | 35606 | No | 1 | 17 | No | 11 | | 64.71 | 0 |
| **Kif3c** | 40886 | Yes | 2 | 18,6 | Yes | 10,2 | | 44 ± 16 | 0 |
| **Kif3c** | 40886 | No | 1 | 7 | Yes | 2 | | 28.6 | 0 |
| **Klp3A** | 40944 | Yes | 1 | 27 | Yes | 16 | 5 cyo adults | 59.3 | 0 |
| **Klp3A** | 40944 | No | 1 | 14 | Yes | 9 | 3 cyo adults | 64.3 | 0 |
| **klp59C** | 35596 | Yes | 1 | 25 | Yes | 23 | | 92 | 0 |
| **klp59C** | 35596 | No | 1 | 16 | No | 13 | | 81.25 | 0 |
| **Ku80** | 27710 | Yes | 1 | 25 | Yes | 7 | | 28 | 0 |
| **Ku80** | 27710 | No | 1 | 25 | No | 24 | | 96 | 0 |
| **Lok** | 35152 | Yes | 1 | 26 | Yes | 20 | 1 with wing spot | 76.92 | 4 |
| **Lok** | 35152 | No | 1 | 25 | No | 24 | 1 with wing spot | 96 | 4 |
| **Map60** | 32458 | Yes | 1 | 11 | Yes | 10 | | 91 | 0 |
| **Map60** | 32458 | No | 1 | 8 | Yes | 8 | | 100 | 0 |
| **Map205** | 32939 | Yes | 6 | 27,27,19,19,25,24 | Yes | 21,10,6,13,14,13 | 5 wing notches | 54 ± 18 | 26 |
| **Map205** | 32939 | No | 6 | 8,15,16,10,21,21 | Yes | 5,12,14,8,21,19 | | 83 ± 13 | 0 |
| **MCPH1** | 38244 | Yes | 1 | 26 | | 9 | | 34.6 | 0 |
| **MCPH1** | 38244 | No | 1 | 15 | | 5 | | 33.3 | 0 |
| **mei-41** | 35371 | Yes | 1 | 27 | Yes | 0 | Lethal after 3$^{rd}$ Instar | . | . |
| **mei-41** | 35371 | No | 1 | 25 | No | 0 | Lethal after 3$^{rd}$ Instar | . | . |
| **Mi-2** | 35398 | Yes | 1 | 25 | Yes | 0 | Lethal after 3$^{rd}$ Instar | . | . |
| **Mi-2** | 35398 | No | 1 | 24 | No | 0 | Lethal after 3$^{rd}$ Instar | . | . |
| **Mi-2** | 33419 | Yes | 2 | 0 | Yes | 0 | Lethal before 3$^{rd}$ Instar | . | . |
| **Mi-2** | 33419 | No | 2 | 0 | No | 0 | Lethal before 3$^{rd}$ Instar | . | . |
| **Mis12** | 38535 | Yes | 1 | 40 | Yes | 0 | Lethal after 3$^{rd}$ instar | . | . |
| **Mis12** | 38535 | No | 1 | 20 | Yes | 0 | Lethal after 3$^{rd}$ instar | . | . |
| **Mis12** | 35471 | Yes | 1 | 12 | Yes | 0 | | 0 | 0 |
| **Mis12** | 35471 | No | 1 | 9 | Yes | 1 | | 11.1 | 0 |
| **Mit(1)15** | 42643 | Yes | 1 | 13 | Yes | 11 | 1 blistered, 1 wing spot | 84.6 | 15 |
| **Mit(1)15** | 42643 | No | 1 | 13 | Yes | 11 | | 84.6 | 0 |
| **mor** | 34919 | Yes | 1 | Dead | . | | Lethal before 3$^{rd}$ instar | . | . |
| **mor** | 34919 | No | 1 | Dead | . | | Lethal before 3$^{rd}$ instar | . | . |
| **mre11** | 39028 | Yes | 1 | 25 | Yes | 0 | | 0 | 0 |
| **mre11** | 39028 | No | 1 | 25 | No | 10 | | 40 | 0 |
| **msps** | 31138 | Yes | 3 | 19,12,25 | Yes | 3,4,1 | 1 wing notch | 18 ± 17 | 5 |
| **msps** | 31138 | No | 2 | 24,6 | No | 20,3 | | 67 ± 24 | 0 |
| **mus209** | 33043 | Yes | 1 | 25 | Yes | 0 | Lethal after 3$^{rd}$ Instar | . | . |
| **mus209** | 33043 | No | 1 | 10 | No | 0 | Lethal after 3$^{rd}$ Instar | . | . |
| **mus309** | 31330 | Yes | 2 | 25,17 | Yes | 9,11 | | 10 ± 1 | 0 |
| **mus309** | 31330 | No | 1 | 22 | No | 21 | | 95.45 | 0 |

| Gene (RNAi) | Stock # | I-CreI | Number of experiments | Number of Larvae per experiment | Heat Shock (1.5 hours) | Normal Adults per experiment | Mutant Phenotype per experiment | % Survival (Avg ± St Dev) | Mutant Wing Phenotype (%) |
|---|---|---|---|---|---|---|---|---|---|
| **mus312** | 34873 | Yes | 1 | 25 | Yes | 0 | | 0 | 0 |
| **mus312** | 34873 | No | 1 | 25 | No | 25 | | 100 | 0 |
| **Nap1** | 35445 | Yes | 1 | 25 | Yes | 20 | 1 with wing spot | 80 | 4 |
| **Nap1** | 35445 | No | 1 | 24 | No | 14 | | 58.33 | 0 |
| **neb** | 28897 | Yes | 1 | 20 | Yes | 13 | | 65 | 0 |
| **neb** | 28897 | No | 1 | 10 | Yes | 9 | | 90 | 0 |
| **Nlp1** | 33688 | Yes | 2 | 29,25 | Yes | 1,0 | | 2 ± 2 | 0 |
| **Nlp1** | 33688 | No | 1 | 6 | Yes | 0 | | 0 | 0 |
| **NudE** | 38959 | Yes | 1 | 17 | Yes | 0 | Lethal after 3rd Instar | . | . |
| **NudE** | 38959 | No | 1 | 12 | Yes | 0 | Lethal after 3rd Instar | . | . |
| **Nuf2** | 35599 | Yes | 1 | 13 | Yes | 7 | | 53.8 | 0 |
| **Nuf2** | 35599 | No | 1 | 8 | Yes | 7 | | 87.5 | 0 |
| **Nuf2** | 36725 | Yes | 1 | 28 | Yes | 0 | Lethal after 3rd Instar | . | . |
| **Nuf2** | 36725 | No | 1 | 25 | Yes | 0 | Lethal after 3rd Instar | . | . |
| **Nurf-38** | 31341 | . | | 0 | . | 0 | Lethal before 3rd instar | . | . |
| **Nurf-38** | 31341 | . | | 0 | . | 0 | Lethal before 3rd instar | . | . |
| **Nurf-38** | 35444 | Yes | | 20 | Yes | 0 | | 0 | 0 |
| **Nurf-38** | 35444 | No | | 21 | No | 2 | | 9.52 | 0 |
| **okr** | 31047 | Yes | 1 | 25 | Yes | 12 | 4 with wing notches | 48 | 16 |
| **okr** | 31047 | No | 1 | 15 | No | 10 | 5 with wing notches | 66.67 | 33 |
| **par-1** | 32410 | Yes | 1 | 0 | Yes | 0 | Lethal before 3rd Instar | . | . |
| **par-1** | 32410 | No | 1 | 0 | No | 0 | Lethal before 3rd Instar | . | . |
| **par-6** | 35000 | Yes | 1 | 25 | Yes | 6 | | 24 | 0 |
| **par-6** | 35000 | No | 1 | 15 | No | 3 | | 20 | 0 |
| **pds5** | 35632 | Yes | 1 | 19 | Yes | 12 | | 63.16 | 0 |
| **pds5** | 35632 | No | 1 | 20 | No | 16 | | 80 | 0 |
| **Pbl** | 36841 | Yes | 1 | 16 | Yes | 0 | Lethal after 3rd instar | . | . |
| **Pbl** | 36841 | No | 1 | 16 | Yes | 0 | Lethal after 3rd instar | . | . |
| **Pbl** | 28343 | Yes | 2 | 13,28 | Yes | 0,0 | Lethal after 3rd instar | . | . |
| **Pbl** | 28343 | No | 2 | 19,23 | Yes | 0,0 | Lethal after 3rd instar | . | . |
| **polo** | 33042 | Yes | 1 | 12 | Yes | 0 | Lethal after 3rd Instar | . | . |
| **polo** | 33042 | No | 1 | 11 | No | 0 | Lethal after 3rd Instar | . | . |
| **Pp4-19C** | 27726 | Yes | 1 | 19 | Yes | 0 | Lethal after 3rd instar | . | . |
| **Pp4-19C** | 27726 | No | 1 | 26 | No | 0 | Lethal after 3rd instar | . | . |
| **Pp4-19C** | 38372 | Yes | 1 | 25 | Yes | 0 | Lethal after 3rd instar | . | . |
| **Pp4-19C** | 38372 | No | 1 | 20 | No | 0 | Lethal after 3rd instar | . | . |
| **PP113C** | 32465 | Yes | 1 | 16 | Yes | 0 | Lethal after 3rd instar | . | . |
| **PP113C** | 32465 | No | 1 | 12 | No | 0 | Lethal after 3rd instar | . | . |
| **PP1-87B** | 32414 | . | 1 | 0 | . | | Lethal prior to third instar | . | . |
| **PP1-87B** | 32414 | . | 1 | 0 | . | | Lethal prior to third instar | . | . |
| **Rpd3** | 34846 | Yes | 1 | 25 | Yes | 0 | Lethal after 3rd Instar | . | . |
| **Rpd3** | 34846 | No | 1 | 25 | No | 0 | Lethal after 3rd Instar | . | . |
| **Rpd3** | 31616 | Yes | 1 | 25 | Yes | 0 | Lethal after 3rd Instar | . | . |
| **Rpd3** | 31616 | No | 1 | 25 | No | 0 | Lethal after 3rd Instar | . | . |
| **Rtf1** | 31718 | Yes | 1 | 25 | Yes | 0 | Lethal after 3rd Instar | . | . |
| **Rtf1** | 31718 | No | 1 | 12 | No | 0 | Lethal after 3rd Instar | . | . |
| **scrib** | 35748 | . | 1 | 0 | . | | Lethal before 3rd instar | . | . |
| **scrib** | 35748 | . | 1 | 0 | . | | Lethal before 3rd instar | . | . |
| **Sin3A** | 32368 | Yes | 1 | 25 | Yes | 2 | | 8 | 0 |
| **Sin3A** | 32368 | No | 1 | 25 | No | 6 | | 24 | 0 |
| **Sir2** | 32481 | Yes | 1 | 25 | Yes | 16 | 3 Wing Notch, 1 Wing Spot | 64 | 16 |
| **Sir2** | 32481 | No | 1 | 16 | No | 15 | | 93.75 | 0 |
| **Sir2** | 31636 | Yes | 1 | 25 | Yes | 15 | 1 Wing Notch | 60 | 4 |
| **Sir2** | 31636 | No | 1 | 29 | No | 29 | | 100 | 0 |
| **SkpA** | 28979 | Yes | 1 | 5 | Yes | 0 | Lethal after 3rd Instar | . | . |
| **SkpA** | 28979 | No | 1 | 4 | Yes | 0 | Lethal after 3rd Instar | . | . |
| **SkpA** | 32870 | Yes | 1 | 0 | . | | Lethal before 3rd instar | . | . |
| **SkpA** | 32870 | No | 1 | 0 | . | | Lethal before 3rd instar | . | . |
| **SMC1** | 34351 | Yes | 1 | 25 | Yes | 24 | | 96 | 0 |
| **SMC1** | 34351 | No | 1 | 11 | No | 11 | | 100 | 0 |
| **SMC2** | 32369 | . | | 0 | . | 0 | Lethal before 3rd instar | . | . |
| **SMC2** | 32369 | . | | 0 | . | 0 | Lethal before 3rd instar | . | . |
| **SNF1A** | 32371 | Yes | 1 | 25 | Yes | 24 | | 96 | 0 |
| **SNF1A** | 32371 | No | 1 | 20 | No | 20 | | 100 | 0 |
| **Spc105R** | 35466 | Yes | 1 | 22 | Yes | 0 | All dead in pupae case | . | . |
| **Spc105R** | 35466 | No | 1 | 22 | Yes | 0 | All dead in pupae case | . | . |
| **Spd-2** | 36624 | Yes | 1 | 25 | Yes | 0 | Lethal before 3rd instar | . | . |
| **Spd-2** | 36624 | No | 1 | 25 | No | 0 | Lethal before 3rd instar | . | . |
| **Spn-A** | 31199 | Yes | 4 | 11,14,7,25 | Yes | 0,1,0,0 | | 2 ± 4 | 0 |
| **Spn-A** | 31199 | No | 5 | 22,21,17,7,32 | Yes | 18,21,10,5,32 | | 82 ± 18 | 0 |
| **spt4** | 31194 | Yes | 1 | 25 | Yes | 6 | | 24 | 0 |
| **spt4** | 31194 | No | 1 | 25 | No | 11 | | 44 | 0 |
| **spt5** | 34837 | . | | 0 | . | | Lethal before 3rd instar | . | . |
| **spt5** | 34837 | . | | 0 | . | | Lethal before 3rd instar | . | . |
| **spt6** | 32373 | . | | 0 | . | 0 | Lethal before 3rd instar | . | . |
| **spt6** | 32373 | . | | 0 | . | 0 | Lethal before 3rd instar | . | . |
| **sub** | 28570 | Yes | 1 | 25 | Yes | 24 | | 96 | 0 |
| **sub** | 28570 | No | 1 | 19 | No | 19 | | 100 | 0 |
| **Su(var)3-9** | 31619 | Yes | 1 | 0 | Yes | 0 | Lethal before 3rd instar | . | . |
| **Su(var)3-9** | 31619 | No | 1 | 0 | No | 0 | Lethal before 3rd instar | . | . |
| **Su(var)3-9** | 32914 | Yes | 1 | 0 | Yes | 0 | Lethal before 3rd instar | . | . |

| Gene (RNAi) | Stock # | I-CreI | Number of experiments | Number of Larvae per experiment | Heat Shock (1.5 hours) | Normal Adults per experiment | Mutant Phenotype per experiment | % Survival (Avg ± St Dev) | Mutant Wing Phenotype (%) |
|---|---|---|---|---|---|---|---|---|---|
| **Su(var)3-9** | 32914 | No | 1 | 0 | No | 0 | Lethal before 3rd instar | . | . |
| **Su(var)3-9** | 33401 | Yes | 1 | 0 | Yes | | Lethal before 3rd instar | . | . |
| **Su(var)3-9** | 33401 | No | 1 | 0 | No | | Lethal before 3rd instar | . | . |
| **Tip60** | 28563 | Yes | 1 | 31 | Yes | 0 | Lethal after 3rd Instar | . | . |
| **Tip60** | 28563 | No | 1 | 25 | No | 0 | Lethal after 3rd Instar | . | . |
| **tefu** | 31635 | Yes | 1 | 30 | Yes | 13 | 1 Wing Spot, 1 Wing Notch | 43.33 | 7 |
| **tefu** | 31635 | No | 1 | 25 | No | 25 | | 100 | 0 |
| **Top2** | 35416 | Yes | 1 | 25 | Yes | 0 | Lethal after 3rd Instar | . | 0 |
| **Top2** | 35416 | No | 1 | 25 | No | 0 | Lethal after 3rd Instar | . | 0 |
| **Top2** | 31342 | Yes | 1 | 19 | Yes | 0 | Lethal after 3rd Instar | . | . |
| **Top2** | 31342 | No | 1 | 12 | Yes | 0 | Lethal after 3rd Instar | . | . |
| **tou** | 31637 | Yes | 1 | 25 | Yes | 17 | | 68 | 0 |
| **tou** | 31637 | No | 1 | 24 | No | 22 | | 91.67 | 0 |
| **tou** | 35790 | Yes | 1 | 25 | Yes | 18 | 1 Wing Spot, 1 Wing Notch | 72 | 8 |
| **tou** | 35790 | No | 1 | 22 | No | 21 | | 95.45 | 0 |
| **tws** | 28714 | Yes | 2 | 4,14 | Yes | 0,0 | Lethal after 3rd Instar | . | . |
| **tws** | 28714 | No | 2 | 8,17 | Yes | 0,0 | Lethal after 3rd Instar | . | . |
| **tws** | 36689 | Yes | 1 | 17 | Yes | 16 | | 94.1 | 0 |
| **tws** | 36689 | No | 1 | 7 | Yes | 7 | | 100 | 0 |
| **vtd (rad21)** | 36786 | Yes | 1 | 25 | Yes | | Lethal after 3rd Instar | . | . |
| **vtd (rad21)** | 36786 | No | 1 | 20 | No | | Lethal after 3rd Instar | . | . |
| **woc** | 27057 | Yes | 1 | 25 | Yes | 0 | | 0 | 0 |
| **woc** | 27057 | No | 1 | 30 | No | 1 | | 3.33 | 0 |
| **Xnp** | 29444 | Yes | 1 | 25 | Yes | 0 | Lethal after 3rd instar | . | . |
| **Xnp** | 29444 | No | 1 | 11 | No | 0 | Lethal after 3rd instar | . | . |
| **Xnp** | 32894 | Yes | 1 | 11 | Yes | 10 | | 90.91 | 0 |
| **Xnp** | 32894 | No | 1 | 22 | No | 16 | | 72.73 | 0 |

**Table S1: Synthetic lethality screen**
117 RNAi gene knockdowns were screened for synthetic lethal interaction upon induction of acentric chromosome formation. Percent survival was determined by taking the average number of larvae that eclosed into viable adult flies per experiment.

| Mode of separation | I-CreI Alone | I-CreI, *nod*$^4$ |
|---|---|---|
| **Sliding** | 48% | 42% |
| **Unzipping** | 22% | 25% |
| **Dissociating** | 13% | 16% |
| **Fails to separate** | 17% | 17% |
| | **N = 19** | **N = 12** |

**Table S2: *nod* loss-of-function mutant does not disrupt the accuracy of acentric sister separation**
Nod does not influence the frequencies of the three modes of acentric separation. Modes by which acentric sister chromatids separate in control cells or in *nod* mutant background. Acentric sister chromatids fail to separate, separate by sliding laterally past one another, or by unzipping from one another. Data for this table is found in Karg *et al*., 2017. All values are not statistically significant (P>0.05, two-sided t-test).
