## Supplemental Movies Description for "Kinetochore-independent mechanisms of sister chromosome separation": Supplementary Movies.pdf

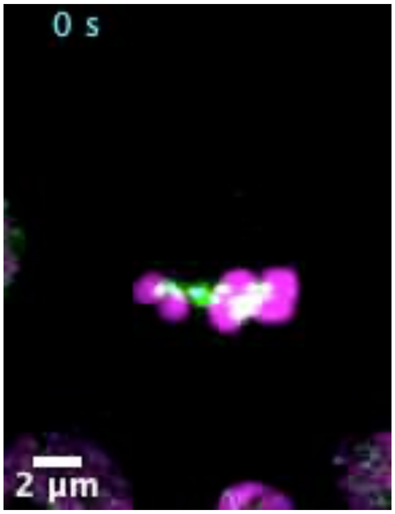


**Movie S1:** Third instar neuroblast mitotic division with I-CreI-induced acentrics and GFP-tagged cohesin. Chromosomes are labeled with H2Av histone variant tagged with RFP (magenta) and cohesin is labeled with Rad21 tagged with GFP (green). Time lapse: 5 seconds. 7 frames per second. Scale bar: 2 μm. This movie corresponds to Figure 2B.

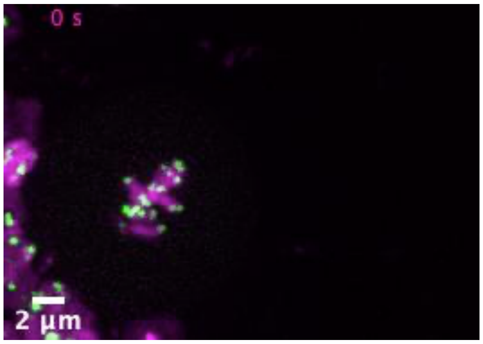


**Movie S2:** Third instar neuroblast mitotic division with I-CreI-induced acentrics and GFP-tagged telomeres. Chromosomes are labeled with H2Av histone variant tagged with RFP (magenta) and telomeres are labeled with HOAP tagged with GFP (green). Time lapse: 18 seconds. 7 frames per second. Scale bar: 2 μm. This movie corresponds to Figure 3A.

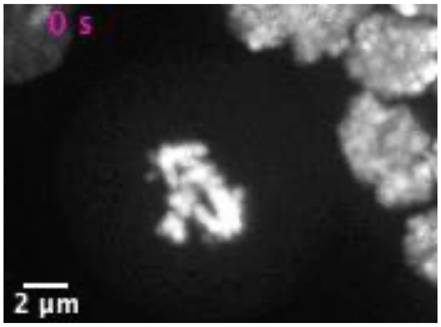


**Movie S3:** Third instar neuroblast mitotic division with I-CreI-induced acentrics and RNAi knockdown of EB1. Chromosomes are labeled with H2Av histone variant tagged with RFP (white). Time lapse: 8 seconds. 7 frames per second. Scale bar: 2 μm. This movie corresponds to Figure 4B.

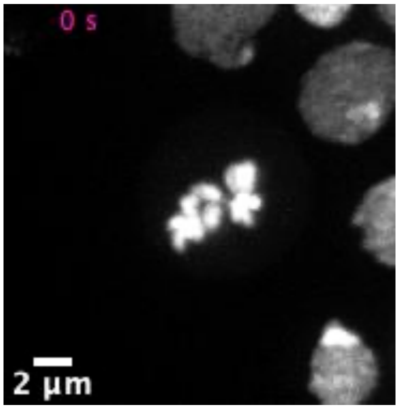


**Movie S4:** Third instar neuroblast mitotic division with I-CreI-induced acentrics and RNAi knockdown of Topoisomerase II. Chromosomes are labeled with H2Av histone variant tagged with RFP (white). Time lapse: 10 seconds. 7 frames per second. Scale bar: 2 μm. This movie corresponds to Figure 6B.

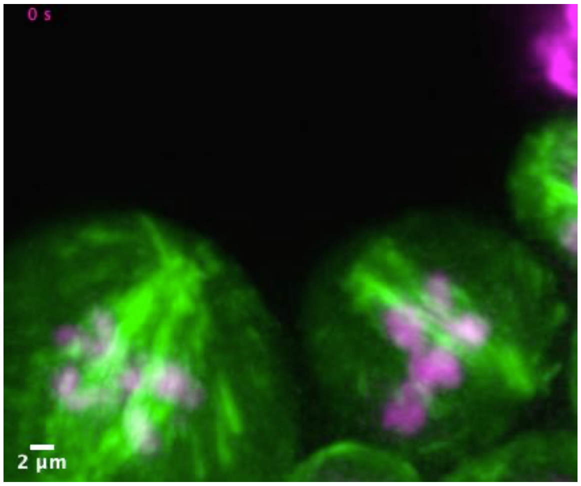


**Movie S5:** Third instar neuroblast mitotic division with I-CreI-induced acentrics and GFP-tagged EB1. Chromosomes are labeled with H2Av histone variant tagged with RFP (magenta) and EB1 is labeled with EB1 tagged with GFP (green). Time lapse: 5 seconds. 7 frames per second. Scale bar: 2 µm. This movie corresponds to Figure 7B.
