## Supplementary figures and images for "Kinetochore-independent mechanisms of sister chromosome separation"

### Movie S1 Acentrics with cohesin-GFP.tif

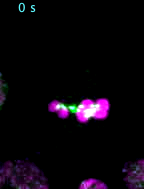

### Movie S2 Acentrics with telomere-GFP.tif

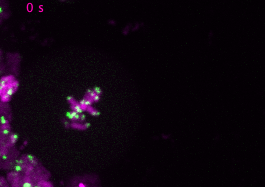

### Movie S3 Acentrics with EB1 RNAi.tif

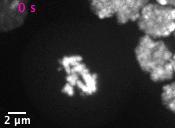

### Movie S4 Acentrics with Topo II RNAi.tif

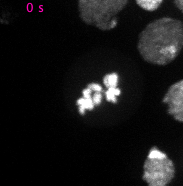

### Movie S5 Acentrics with EB1-GFP.tif

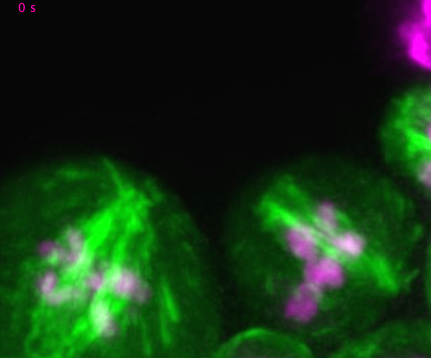
